# iPSC Neurodegenerative Disease Initiative isogenic CAG repeat iPSC line for Huntington’s disease

**DOI:** 10.64898/2026.06.30.735662

**Authors:** Lisa Salazar, Mara S. Burns, Jennifer T. Stocksdale, Keona Q. Wang, Gloria Cao, Ricardo Miramontes, Nicolette R. McClure, Leanne Ho, Amber Rose Keith, Margaret Sutherland, Mark R Cookson, Michael Ward, William C. Skarnes, Leslie M Thompson

## Abstract

**Purpose of Research:** The generation of iPSC lines expressing 21, 56 and 79 glutamine repeats within the HTT protein and homozygous KO of *HTT* in the KOLF2.1J background as an additional disease series within the iPSC Neurodegenerative Disease Initiative (iNDI) collection.

**Major Findings:** All iPSCs, even those expressing long repeats of 79Q or HTT KO, were capable of differentiating to striatal and cortical neurons, astrocytes and microglia using established protocols. General quality control stains and morphological analyses are described for each differentiation. A selected set of assays were carried out on differentiated cells; expanded repeat expressing astrocytes showed altered expression of astrocyte protein markers and morphological characteristics, and striatal neurons showed altered DARPP-32/CTIP2 colocalization. mRNAseq carried out for striatal neurons showed high similarities in gene expression changes between 79Q and KO lines compared to the unexpanded repeat.

**Conclusions:** The KOLF2.1J isogenic CAG repeat series serves as a community resource to study HD mechanisms with the potential for direct comparison across other neurodegenerative diseases through the iNDI collection.

## INTRODUCTION

Huntington’s disease (HD) is an inherited neurodegenerative disorder characterized by motor, psychiatric and cognitive symptoms that is caused by a CAG repeat expansion in the first exon of the huntingtin gene (HTT)^1^ and the length of the CAG tract partially accounts for the age of symptom onset with an additional significant contribution from modifier genes including those in the mismatch repair pathways^2–4^. Although HTT is ubiquitously expressed, striatal and cortical tissues are most affected, with medium spiny neurons (MSNs) particularly vulnerable^5^.

The advent of induced pluripotent stem cells (iPSCs) two decades ago^6^ has provided the ability to study the effects of the CAG repeat expansion in an endogenous patient-derived background within a myriad of cell types. However, differences in genetic background between patients can strongly influence results, often making it a challenge to conclusively attribute differences observed between HD and control cells to the presence of the mutation without the use of large numbers of lines. In more recent years, genome editing technologies have been utilized to create human isogenic iPSC lines, including isogenic pairs generated by the gene correction of HD lines^7, 8, 9^ or by the introduction of an expanded CAG tract into the unexpanded HTT locus of control lines.^10^ Isogenic series made by the introduction of multiple expanded CAG repeat lengths have also been derived^11, 12^.

Among allelic series carrying a broad range of CAG repeat lengths engineered into a normal repeat background is the RUES2 female embryonic stem cell (ESC) series whereby CRISPR technology was used to introduce repeat lengths of 45Q, 50Q, 58Q, 67Q, and 74Q and homozygous and heterozygous HTT knockout.^12^ These lines recapitulate previously observed developmental abnormalities with highly expanded repeats, including increased numbers of progenitor cells, aberrant rosette formation, and altered gene expression in developmental pathways, as well as novel chromosomal instability.^12, 13^ The same phenotypes were observed in homozygous HTT knockout lines. These lines have been used by several groups to identify HD phenotypes in 2D and organoid cultures.^14, 15^ A second allelic series, the IsoHD series, was generated using TALENs to introduce expanded CAG repeats into female H9 ESCs, producing lines with 30Q, 45Q, 65Q, and 81Q.^11^ These lines showed CAG length-associated abnormalities in mitochondrial respiration and oxidative stress as well as an increased sensitivity to DNA damage in differentiated neural cells. Microglia derived from these lines were found to be hyper-reactive to stimulation, released elevated levels of reactive oxygen species, and were more susceptible to exogenous stress and apoptosis.^16^

While the generation of isogenic lines have provided critical information relating to HD mechanisms, they were each generated in different genetic backgrounds and do not provide the opportunity to directly compare to other neurodegeneration disease-causing mutations. The iPSC Neurodegenerative Disease Initiative^17^ is a collaborative effort from the Center for Alzheimer’s and Related Dementias (NIH), with support from the Chan Zuckerberg Initiative, Aligning Science Across Parkinson’s (ASAP), and The Jackson Laboratory to create a library of human induced pluripotent stem cells (iPSCs) for Alzheimer’s and Related Dementias (ADRDs), ALS, and other neurodegenerative disorders. Using CRISPR/Cas9-based genetic engineering, more than 100 variants associated with neurodegenerative disorders have been introduced into control KOLF2.1J iPSCs, a well-characterized male reference cell line that is genetically stable upon CRISPR/Cas9 editing and can differentiate along commonly used lineages.^18^ Use of these isogenic lines allows the study of how each variant contributes to disease in a genetically defined background and enables comparisons across neurodegenerative disorders. This library of isogenic cell lines is a community resource available through The Jackson Laboratory (https://www.jax.org/jax-mice-and-services/ipsc).

Here we describe the generation of *HTT* iPSC lines with 21, 56 and 79 polyglutamine repeats, hereafter designated as 21, 56 and 79Q, and homozygous KO of HTT in the KOLF2.1J background as an additional disease series for the iNDI collection. All iPSCs, even those with long repeats of 79Q and KO, are capable of differentiating to striatal and cortical neurons, astrocytes and microglia using established protocols. General quality control stains, morphological analysis and synaptic co-staining are described. We observed CAG repeat length-dependent changes in GLAST expression and morphological characteristics of astrocytes, as well as altered DARRP-32/CTIP2 colocalization of KO striatal neurons. Microglia and cortical neurons did not display overt functional differences in the assays tested, and no changes were observed in synaptic co-staining for PSD95/SYP, suggesting the repeat expansions in the wild type KOLF2.1J background elicit subtle changes. Transcriptomic analysis of striatal differentiations, including HTT isogenic and knockout lines, was also performed and the most significant gene expression differences were observed in the 79Q and HTT knockout lines. Differential gene expression analysis highlights dysregulated pathways observed in HD models, including those related to CREB signaling, cell adhesion, and cAMP-mediated signaling, although the specific DEGs contributing to these pathways differed across lines. These lines can serve as a community resource to study HD mechanisms with the potential for direct comparison across neurodegenerative diseases.

## METHODS

### Generation of a HTT CAG allelic series in iPSCs

The KOLF2.1J reference cell line,^18^ which is the reference clonal cell line for the iNDI consortium^17^ and contains 18 and 20 uninterrupted CAGs followed by a CAACAG that encodes 20 and 22 uninterrupted glutamines (Q), was used to create an allele-specific CAG isogenic series using CRISPR/Cas9 technology (Table S1). We first introduced a heterozygous 3bp deletion immediately upstream of the CAG repeat region, then designed a sgRNA (sgRNA2) to the mutated sequence, thereby targeting the same allele for introduction of each expanded repeat. A ‘base’ homology donor plasmid conceptually similar to previous approaches^12^ was synthesized by GenScript (Piscataway, NJ) to contain HTT exon1 with 19 CAG repeats (19CAG-CAA-CAG; 21Q) and approximately 1kb homology arms (Gene ID: 3064). To create the expanded repeat exon1 donor plasmids with adult and juvenile CAG lengths, the HTT exon 1 region was PCR amplified from genomic DNA obtained from iPSCs (CS03iHD-53n3) or from a commercially available HTT donor plasmid (pJOP-HTT-HR81Q; Addgene plasmid # 92250 deposited by Mahmoud Pouladi^11^), and cloned into the ʻbase’ plasmid using XmnI and BbsI sites which flank the CAG tract. The primers used for the CAG tract amplification were polyCAG_Fw and polyCAG_Rv (Table S1).^12^ CRISPR/Cas9 editing for the expanded repeats proceeded in two steps.^19^ In the first step, sgRNA1+Cas9 RNP introduced a heterozygous 3bp deletion immediately upstream of the CAG tract, allowing for allele-specific introduction of desired length CAG repeats. In the second step, sgRNA2, targeting the 3bp deletion allele was introduced along with Cas9 protein and HTT or mHTT exon1 plasmid donors. Four knock-in clones for each CAG length were confirmed by PCR amplification and Sanger sequencing using PCR primers HTT_PF3 and HTT_PR3 with sequencing primer HTT_SF1 (Table S1). One clone for each CAG length was further validated by nanopore sequencing of the expanded repeat, confirming expanded repeats of 54CAG-CAA-CAG (56Q) and 77CAG-CAA-CAG (79Q). Sanger sequencing indicated an insertion of 19CAG-CAA-CAG for the 21Q line and a repeat of 18CAG-CAA-CAG (20Q) for the unedited allele of all lines. Subcloning and sequencing confirmed clonality, and Western blot validated expression of full length HTT and mHTT. Genomic integrity was confirmed by SNP Array.

### Generation of homozygous HTT Knockout iPSC lines

KOLF2.1J cells were nucleofected with Cas9 protein, sgRNA3, sgRNA4, and donor plasmid to facilitate excision of the entire HTT coding region. However, PCR amplification using primers across the cut sites at the 5’ (HTT_KO_PF1; HTT_KO_INT_PR1) and 3’ (HTT_KO_INT_PF1; HTT_KO_PR1) ends or deletion junction (HTT_KO_PR1; HTT_KO_PR1) indicated that this approach produced only heterozygous knockouts. Therefore, a pool of heterozygous knockout clones was re-targeted with sgRNA5 and sgRNA6 in conjunction with a donor oligo to remove exon 6 of the other allele, thereby producing homozygous HTT knockout lines. Three clones were confirmed to be compound heterozygous KO clones (deletion of CDS/deletion of exon 6) by PCR amplification and Sanger sequencing across the exon 6 target sequence using PCR primers HTT E6 PF1 and HTT E6 PR1 with sequencing primer HTT E6 SR. Two clones *HTT*KO_1, and *HTT*KO_2 were further evaluated by Western blot analysis and confirmed null for expression of HTT protein. Genomic integrity was confirmed by Array Comparative Genomic Hybridization (aCGH). sgRNA, primer, and donor sequences are provided in Table S2.

### qPCR for pluripotency markers

To assess pluripotency, expression levels of *NANOG, OCT4*, and *SOX2* in the 21Q, 56Q, 79Q, *HTT*KO_1, and *HTT*KO_2 iPSC lines were compared to the parental KOLF2.1J line and NGN2-induced 21Q cortical neurons. Total RNA was isolated using the RNeasy Mini Kit (Qiagen; #74106) with QIAshredders (Qiagen; #79656) for homogenization and on-column DNase I treatment (Qiagen; #79254) to remove genomic DNA. RNA purity and concentration were verified using a NanoDrop spectrophotometer by monitoring A260/280 and A260/230 ratios. cDNA was synthesized from 1 µg of total RNA using the QScript™ cDNA SuperMix (QuantaBio; #101414-106) on a Bio-Rad T100 Thermal Cycler. The resulting cDNA was diluted to 5 ng/µL in nuclease-free water and stored at −20 °C. Quantitative PCR (qPCR) was performed on a QuantStudio™ 7 Flex Real-Time PCR System using 20 ng of cDNA per reaction. Technical triplicates were analyzed for each cell line. Relative expression was calculated using ΔCt by subtracting the mean Ct of the housekeeping gene, RPLP0, from the mean Ct of each target gene (*NANOG*, *OCT4*, and *SOX2)*.

### Western blot of for HTT length

Cell pellets were broken in a modified RIPA buffer containing: 10mM Tris, 150mM NaCl, 1mM EDTA, 1% NP40, 0.5% SDS. Phosphatase inhibitors 2 (Millipore Sigma, P5726) (1:1000) and 3 (Millipore Sigma P0044) (1:1000), 1 mM PMSF, 10 μg/mL aprotinin (Sigma-Aldrich A1153), 10 μg/mL leupeptin (Sigma-Aldrich L2884), and one Pierce mini protease pellet (Fisher Scientific A32953) per 10 mL of lysis buffer were added. Lysates were sonicated (3 times for 10 seconds at 40% power) and protein concentration was analyzed via Lowry assay. 50μg of protein was then used for SDS/PAGE. NuPage 3-8% Tris-Acetate Mini Protein Gels (Thermo Fisher Scientific EA03785BOX) were used with Tris-acetate running buffer (Fisher Scientific LA0041). The gel was then transferred onto Immobilon-FL PVDF (Millipore Sigma IPFL00010). Whole protein was visualized using Revert Total Protein Stain assay (LICOR Biosciences 926-11016), and the membrane was blocked with Intercept (TBS) Blocking Buffer (LICOR biosciences 927-60010) for 1 hour. The membrane was then incubated in primary antibody (Abcam ab109115 (Lot# GR3339173-2), 1:1000) overnight, washed three times with TBS-0.1% Tween-20, and incubated for 1 hour in near-infrared-conjugated secondary antibody (LICOR biotech IRDye) in intercept block supplemented with 0.1% Tween-20. Membranes were imaged on a LICOR scanner.

### NGN2-mediated differentiation to cortical neurons

Cortical neuron populations were generated as previously described^18, 20^ with some modifications. Induced pluripotent cell lines were maintained in mTeSR Plus medium (Stem Cell Technologies 100-0276) on hESC-qualified matrigel (Corning) and passaged using ReLeSR (Stem Cell Technologies100-0484) at 80% confluence in the presence of CEPT (Chroman1-Tocris 7163, Emricasan-Seleck Chemicals S7775, Polyamine supplement-Sigma P8483, Trans-ISRIB-R&D Systems-5284.^21^ All three lines (21Q, 56Q, 79Q) were transfected via Nucleofection (LONZA VPH-5022) with the PB-TO-hNGN2 plasmid, a gift from iPSC Neurodegenerative Disease Initiative (iNDI) & Michael Ward (Addgene plasmid # 172115; http://n2t.net/addgene:172115; RRID:Addgene_172115).^18^ Transfected cells were grown for multiple passages in the presence of 200 ng/mL Puromycin (Invivogen ant-pr-1) to select for plasmid expression as determined by BFP (blue fluorescent protein) expression. Once a high percentage of the cell population was found to express BFP, NGN2-iPSCs were dissociated to single cell with Accutase (ThermoFisher NC9464543) and seeded at ∼1x10^6^ cells per hESC-qualified matrigel (Corning) coated 6 well plates to begin cortical differentiation in Induction media: Knockout DMEM/F12 (ThermoFisher); N2 supplement 100X (ThermoFisher); non-essential amino acids 100X (ThermoFisher), and supplemented with Doxycycline at a final concentration of 1μM (Sigma) and CEPT. Induction medium was changed daily for 3 days with the addition of Uridine (U) and Fluorodeoxyuridine (FdU) at 1mM each (Sigma 3750, Sigma 0503) on day 3. On day 4, cells were dissocated to single cell using Accutase and seeded at 1x10^6^ cells per Poly-D-Lysine coated 6 wells (Sigma P6407) and 8x10^4^ cells per coated glass coverslip. Cells were plated into Cortical Neuron Culture Medium 1 (CM1): 1:1 Knockout DMEM/F12: BrainPhys neuronal medium without Phenol-Red (STEMCELL Technologies); B27 supplement, 50X (ThermoFisher); BDNF (10 μg/ml, STEMCELL Technologies ) in PBS containing 0.1% BSA (ThermoFisher); NT-3 (10 μg/ml, Preprotech) in PBS containing 0.1% BSA, GDNF (10 μg/ml, STEMCELL Technologies) in PBS containing 0.1% BSA; laminin final con. 1 μg/ml (ThermoFisher), Doxycycline (1uM), U (1uM), and FdU (1uM). Cells were given a half medium change on day 7 with CM1 then maintained with half media changes every 3-4 days for an additional 21 days in Cortical Neuron Culture Medium 2 (CM2). CM2: BrainPhys neuronal medium without Phenol-Red (STEMCELL Technologies); B27 supplement, 50X (ThermoFisher); BDNF (10 μg/ml, STEMCELL Technologies) in PBS containing 0.1% BSA (ThermoFisher); NT-3 (10 μg/ml, Preprotech) in PBS containing 0.1% BSA, GDNF (10 μg/ml, STEMCELL Technologies) in PBS containing 0.1% BSA; laminin final con. 1 μg/ml (ThermoFisher), Doxycycline (1μM), U (1μM), and FdU (1μM). Cells were fixed in 4% PFA at day 28 for immunofluorescence.

### NGN2-mediated differentiation immunocytochemistry

After 28 days in culture, cortical neuron populations were fixed with 4% paraformaldehyde (Fisher Scientific # 50980487) for 10 minutes at room temperature, then washed three times with PBS (Corning # 21030CV). Cells undergo antigen retrieval with EDTA at 65°C for 10 minutes. Cells were then permeabilized with 0.3% Triton-X (Sigma #T8787) in PBS for 10 minutes and then blocked with 2% goat serum (Thermo Fisher #16210-064), 3% BSA (Thermo Fisher # 15260-037), 0.1% Triton-X, and 0.3M Glycine (Fisher # BP381-1) in PBS for 1 hour at room temperature and then incubated in primary antibody diluted in block, overnight at 4°C (anti-MAP2 (1:1000; Synaptic Systems 188004), anti-TBR1 (1:250; Abcam ab31940), anti-NeuN (1:1000; Millipore ABN90P). Primary antibody was removed, and cells washed three times with PBS and then incubated for 1 hour in secondary antibody diluted 1:1000 in block, in the dark at room temperature (Alexa Fluor Goat IgG (H+L) Secondary Antibody, Thermo Fisher Scientific). Cells were washed with PBS for three times and then washed in PBS containing Hoechst 33342 (Sigma #14533) for 10 minutes and then a final wash in PBS. Coverslips were mounted with Fluoromount-G® (Fisher # OB10001) and allowed to dry.

### NGN2-mediated differentiation Analysis

Images were acquired at 20× magnification using an Olympus FV3000 Confocal Laser Scanning Microscope (n = 4 images per coverslip; 1 differentiation). Cortical neurons were analyzed in Imaris (version 11.0.0). Total nuclei (DAPI) were quantified using the Imaris Spots tool. TBR1- and NeuN-positive cells were identified with the Imaris Surface tool, and surface measurements were used to determine nuclei colocalizing with each marker. Colocalized cells were normalized to the total number of nuclei per image and analyzed in GraphPad Prism.

### Astrocyte Differentiation

Astrocyte differentiation was modified based on previous publication.^22^ KOLF iPSC lines were differentiated into neural progenitor cells (NPCs) according to manufacturer’s instructions (Gibco #A1647801, MAN0008031). CEPT was added at each passage point for 24 hours to increase cell survival. NPCs were cultured to 90–95% confluency, then seeded at 1X10^5^ cells/cm^2^ on hESC-qualified matrigel (Corning) for four passages before induction of astrocyte differentiation. To begin astrocyte differentiation, passage four NPCs were seeded at 7.5x10^4^ cells/cm^2^ in Neural Expansion Medium (Gibco #A1647801) and the following day (day 0), half the medium was replaced with astrocyte differentiation medium (ADM) 1. Half media changes were performed every other day for the remainder of the differentiation. All subsequent passages were performed by washing cells with HBSS without Mg Ca (Gibco) three times, then incubating in StemPro Accutase (Gibco) at 37°C for 8-10 minutes, counting using the Countess II Fluorescence automated cell counter with trypan blue exclusion and seed on hESC-qualified matrigel (Corning) at 6x10^4^ cells/cm^2^ at day 7, day 15, and day 30, 5x10^4^ cells/cm^2^ at day 45 and 4x10^4^ cells/cm^2^ at day 60. On day 7, the cells were replated in ADM1. On day 15, the cells were replated in 50% ADM1, 50% ADM2. Subsequent half media changes were performed using ADM2. On day 30, day 45, and day 60 the cells were seeded in 100% ADM3. iAstrocytes were harvested and banked on day 67. Quality control immunocytochemistry was performed at each passage time point for PSC, NPC, neural, and astrocytic markers: DCX (1:500, Millipore AB2253), GFAP (1:1000, Abcam ab4674), GLAST (1:100, Miltenyi Biotec 130-095-822), MAP2 (1:1000, Synaptic Systems 188004), Nestin (1:1000, Millipore MAB5326), OCT4 (1:500, Millipore MAB4401), PAX6 (1:250, Abcam ab5790), S100b (1:1000, Sigma-Aldrich S2532), SOX2 (1:500, Millipore ab5603), SOX9 (1:500, Sigma-Aldrich ab5535), and TUJ1 (1:1000, Millipore MAB5564).

### Astrocyte Immunocytochemistry

Cells were fixed using 4% paraformaldehyde (Electron Microscopy Sciences) in PBS for 10 minutes at room temperature. Depending on antigen, cells were permeabilized with 0.3% Triton X-100 (Sigma-Aldrich) in PBS for 10 minutes at room temperature. Cells were blocked for 1 hour in 10% goat serum (Gibco), 1% BSA (Gibco), 0.1% Triton X-100 in PBS for 1 hour at room temperature. Cells were incubated with primary antibodies overnight at 4°C and then washed three times with PBS before incubation in 1:1000 Alexa-Fluor 488, 594, 647 secondary antibodies (Invitrogen) for 1 hour at room temperature and subsequently incubated with Hoechst 33342 (Sigma-Aldrich) for 10 minutes at room temperature. Stained slides were mounted with Fluoromount-G (Southern Biotech). Quality control stained iAstrocytes were imaged with a Keyence fluorescence microscope at 10x and 20x. Analysis and visualization of images were performed by Imaris.

### Astrocyte GLAST and SOX9 intensity and Sholl morphometric analysis

Individual iAstrocytes (n = 40 cells per line, 1 biological differentiation replicate per CAG repeat) were imaged on a Fluoview FV3000 Olympus Confocal Microscope at 40× magnification. iAstrocytes were analyzed with Imaris (version 10.2.0) Filaments manual function to determine total filament length using AutoPath and Segment Type by tracing the GLAST channel. The size of the nucleus (DAPI) was set as the starting point and then processes were traced from that point. Additionally, based on the paths, Sholl^23^ on Imaris was conducted to assess the complexity of each iAstrocytes calculated by the number of intersections and concentric circles at 1 μm sequentially distant radii.

### Microglia Differentiations

iPSC-derived microglia (iMG) were differentiated as previously described.^24^ Briefly, iPSCs were differentiated into hematopoietic progenitors by passaging using ReLeSR and trituration onto 6-well Geltrex-coated plates (Thermo, A1569601) in mTeSR1 at a density of 30-50 colonies per well. Once colonies were 100-200 µm (24-48hrs after plating), cells were transferred to Medium A from the STEMdiff Hematopoietic Kit (STEMCELL Technologies, 05310) (day 0) and fed every 2 days. After 3-4 days (day 3 or 4), once colonies reached 500-1000um size, cells were changed to Medium B and fed every 2 days until day 11 or 12 when small round HPCs began to lift into the media. On day 10 or later, media from the cells was gently collected along with non-adherent CD43+ HPCs. Collected HPCs were pelleted and frozen down in BamBanker freezing solution (Wako, NC9582225) for long-term storage. HPCs were thawed at a density of 100,000-200,000 cells per well of a Geltrex-coated 6-well plate into iMG medium freshly supplemented with 100 ng/mL IL-34 (200-34; Peprotech], 50 ng/mL TGFb1 (100-21; Peprotech), and 25 ng/mL M-CSF (300-25; Peprotech). iMG medium: DMEM/F12 (Thermo, 11039021), 2% Insulin-transferrin-selenium (Thermo, 41400045), 1% Glutamax (Thermo, 35050061), 1% NEAA (Thermo, 111140050), 400 μM Monothioglycerol (Sigma, M6145), 5ug/mL Insulin (Sigma, I2643-25mg), 2% B27 (Thermo, 17504001), 0.5% N2 (Thermo, 17502048). Cultures were fed every 48 hours with freshly supplemented iMG media for 28 days and expanded as necessary (maintaining ∼500,000 cells per well). After 25 days in culture, two additional cytokines were added (100 ng/mL CD200 (770006; BioLegend) and 100 ng/mL CX3CL1 (P300-31; Peprotech)) to mature the microglia in a homeostatic brain-like environment.

### Microglia Immunocytochemistry

iMGs were plated onto Vitronectin (Stemcell Technologies, 07180) coated glass coverslips and allowed to acclimate overnight before being fixed with 4% paraformaldehyde (50980487; Fisher Scientific) for 10 minutes at room temperature, then washed three times with PBS (21030CV; Corning). Cells were permeabilized with 0.3% Triton-X (T8787; Sigma) in PBS for 10 minutes and then blocked with 2% goat serum (16210-064; Thermo Scientific), 3% BSA (15260-037; Thermo Scientific), 0.1% Triton-X in PBS for 1 hour at room temperature and then incubated in primary antibody diluted in block (1:200 PU.1 Cell Signaling, 2266s; 1:400 P2RY12 Sigma, HPA014518; 1:1000 CX3CR1 Bio-Rad, AHP1589; 1:1000 IBA1 Fuji film, 019-19841), overnight at 4°C. Primary antibodies were removed, and cells washed three times with PBS and then incubated for 1 hour in secondary antibody diluted 1:1000 in block, in the dark at room temperature (Alexa Fluor Goat IgG (H+L) Secondary Antibody, Thermo Scientific). Cells were washed with PBS for three times and then washed in PBS containing Hoechst 33342 (Sigma #14533) for 10 minutes and then a final wash in PBS. Coverslips were mounted with Fluoromount-G® (Fisher # OB10001) and allowed to dry. Images were acquired with a Keyence BZ-X810 Widefield Microscope with a 20X objective.

### Microglia Phagocytosis Assay

Phagocytosis was measured using a previously described protocol.^24^ iMGs were plated at 33,000 cells per well of a Matrigel coated 96-well plate in conditioned media from their culture wells and allowed to acclimate overnight. The next day, iMGs were imaged once using the IncuCyte S3 Live-Cell Analysis System (Sartorius) before the addition of *S. aureus* particles. Then pHRodo Green dyed *S. aureus* particles (Thermo, P35367) were diluted into iMG media with fresh cytokines at a density of 20,000 particles per well and added to the 96-well iMG plate. Images of phase and fluorescence for 4 fields of view in each of four wells were captured for each condition every 30 minutes for 24 hrs. Using IncuCyte 2019B software (Sartorius), image masks for fluorescent signal (phagocytosis of each substrate) were normalized to cellular confluence (brightfield). GraphPad (Prism) was used to analyze differences in normalized fluorescence by two-way ANOVA and Tukey’s multiple comparisons corrections.

### Striatal Neuron Differentiation

Neuronal differentiation was performed once iPSC colonies reached 60–70% confluency as previously described.^25^ Differentiation was initiated (Day 0) by washing iPSC colonies with phosphate-buffered saline pH 7.4 (PBS – Gibco) and switching to SLI medium (Advanced DMEM/F12 (1:1) supplemented with 2 mM Glutamax^TM^ (Gibco), 2% B27 without vitamin A (Life Technologies), 10 µM SB431542, 1 µM LDN 193189 (both Stem Cell Technologies), 1.5 µM IWR1 (Tocris)) with daily medium changes. Four days later, cells were pretreated with 50 nM Chroman 1, washed with PBS, and passaged 1:2 with StemPro Accutase (Invitrogen) for 5 minutes at 37 °C. Cells were replated onto plates coated with hESC qualified matrigel (1 h at 37 °C) in SLI medium containing CEPT cocktail for 1 day after plating and continued daily feeding with SLI medium. At day 8, cells were passaged 1:2 as above and replated in LIA medium (Advanced DMEM/F12 (1:1) supplemented with 2 mM Glutamax^TM^, 2% B27 without vitamin A, 0.2 µM LDN 193189, 1.5 µM IWR1, 20 ng/ml Activin A (Peprotech)) with CEPT cocktail for 1 day after plating and daily feeding was continued through day 16. At day 16, cells were plated in 6-well Nunclon plates coated with PDL and hESC Matrigel at 1 × 10^6^ per well in SCM1 medium (Advanced DMEM/F12 (1:1) supplemented with 2 mM Glutamax^TM^, 2% B27 (Invitrogen), 10 µM DAPT, 10 µM Forskolin, 300 µM GABA, 3 µM CHIR 99021, 2 µM PD 0332991 (all Tocris), 1.8 mM CaCl_2_, 200 µM ascorbic acid (Sigma-Aldrich), 10 ng/ml BDNF (Peprotech)). Medium was 50% changed every 2-3 days. On day 23, medium was fully changed to SCM2 medium (Advanced DMEM/F12 (1:1): Neurobasal A (Gibco) (50:50) supplemented with 2 mM Glutamax^TM^), 2% B27, 1.8 mM CaCl_2_, 3 µM CHIR 99021, 2 µM PD 0332991, 200 µM ascorbic acid, 10 ng/ml BDNF) and 50% medium changes performed every 2–3 days. Cells were considered mature at day 37.

### Striatal neuron immunocytochemistry and image analysis

At day 37, striatal neurons were fixed with 4% paraformaldehyde (Fisher Scientific # 50980487) for 10 minutes at room temperature, then washed three times with PBS (Corning # 21030CV). Cells were permeabilized with 0.3% Triton-X (Sigma #T8787) in PBS for 10 minutes and then blocked with 2% goat serum (Thermo Fisher #16210-064), 3% BSA (Thermo Fisher # 15260-037), 0.1% Triton-X, and 0.3M Glycine (Fisher # BP381-1) in PBS for 1 hour at room temperature and then incubated in primary antibody diluted in block, overnight at 4°C (anti-DARPP32 (1:200 Abcam 40801), anti-CTIP2 (1:500 Abcam 18465), anti-FOXP1 (1:1000 Abcam 16645), anti-MAP2 (1:1000 Synaptic Systems 188004), anti-NEUN (1:1000, Millipore Sigma ABN90P), anti-synaptophysin (1:200 Abcam 8049), anti-PSD95 (1:100, Abcam 18258)). After removing primary antibody, cells were washed three times with PBS and then incubated for 1 hour in secondary antibody diluted 1:1000 in block, in the dark at room temperature (Alexa Fluor 488, 555, 647 Goat IgG (H+L) Secondary Antibody, Thermo Fisher Scientific). Cells were washed with PBS three times and then placed in PBS containing Hoechst 33342 (Sigma #14533) for 10 minutes, followed by a final wash in PBS. Coverslips were mounted with Fluoromount-G® (Fisher # OB10001) then imaged on a Fluoview FV3000 Olympus Confocal Microscope at 20x and 40x. Three random images were taken per coverslip. DARPP-32, CTIP2, NEUN, and total nuclei (DAPI) were quantified using Imaris spots tool (Imaris Single Full software, BITPLANE). Imaris colocalization tool was used to count the number of DARPP-32+/CTIP2+/DAPI+ cells and the number of NEUN+/DAPI+ cells. For synaptophysin and PSD95, four random images were taken per coverslip. Imaris surface and spot tools were used to identify synapses, and quantification was done by the Imaris colocalization tool using a threshold setting of 1.25. Data was analyzed with GraphPad Prism software using one-way ANOVA assuming a normal distribution.

### mRNA Sequencing

RNA from Day 37 cell pellets was extracted using the RNeasy mini kit with DNaseI for gDNA elimination (Qiagen, 74106, 79254). All samples had an RNA integrity number (RIN) of 9.6 or greater. rRNAs were removed and libraries generated using TruSeq Stranded mRNA library prep kit with Ribo-Zero (Qiagen). RNA-seq libraries were normalized according to size (Agilent Bioanalyzer 2100 High Sensitivity chip) and sequenced on the NovaSeq X Plus with sufficient paired end (PE) sequencing cycles to obtain 50 million PE reads per sample. Sequenced reads were trimmed for adaptor sequence or low-quality sequence using Trimmomatic^26^ and aligned using HISAT2 using an index created from Ensembl GRCh38 release 104. Gene counts were quantified using featureCounts^27^ and differential analysis was performed DESeq2.^28^ Additional QC metrics were obtained using SAMtools. Ingenuity Pathway Analysis of DEGs (padj < 0.05, FC > 0.6) was used to investigate biological changes in iMSNs, using the entire human genome as the background.^29^

## RESULTS

### Generation of a HTT CAG allelic series in iPSCs

To model the effects of expanded CAG repeat alleles within the specifications of the iNDI project,^17^ we generated a *HTT* allelic series in the KOLF2.1J iPSC line and performed differentiations to disease-relevant cell types including neurons, astrocytes and microglia (Figure 1A). Allele-specific introduction of 56Q and 79Q repeats at the HTT locus used a two-step editing process (Figure 1B). First, CRISPR/Cas9 technology was used in conjunction with a sgRNA (sgRNA1) targeting the sequence immediately upstream of the CAG repeat region in the KOLF2.1J line, which contains 19/21 CAG (21/23Q) repeats. A clone was isolated containing a 3bp deletion in only one of the *HTT* alleles. The deletion was contained within the sgRNA target sequence, leaving the parental CAG tract unaffected, and allowing for a new sgRNA (sgRNA2) to be designed, targeting the 3bp deletion. This allowed for allele-specific insertion of normal (19CAG/21Q) and expanded (54CAG/56Q and 77CAG/79Q) repeats. The 19CAG/21Q was included to ensure that all lines went through the same editing process and that gene-editing itself did not affect any readouts. Four independent iPSC clones for each genotype were evaluated by Sanger sequencing across the targeted region and one clone each of 21Q, 56Q, and 79Q was chosen for full characterization. Expanded alleles were amplified and sequenced to confirm repeat length (Figure S1). Normal and unedited alleles were confirmed by Sanger sequencing, indicating a repeat length of 18CAG/20Q for the unedited allele in all lines (data not shown). Western blot confirmed protein expression of HTT and mHTT (Figure 1C, S2A-C). All lines maintained pluripotency and parental karyotype^30, 31^, evidenced by qPCR for standard markers (Figure S2D) and SNP analysis (Figure S3, Table S3), respectively.

**Figure 1:**
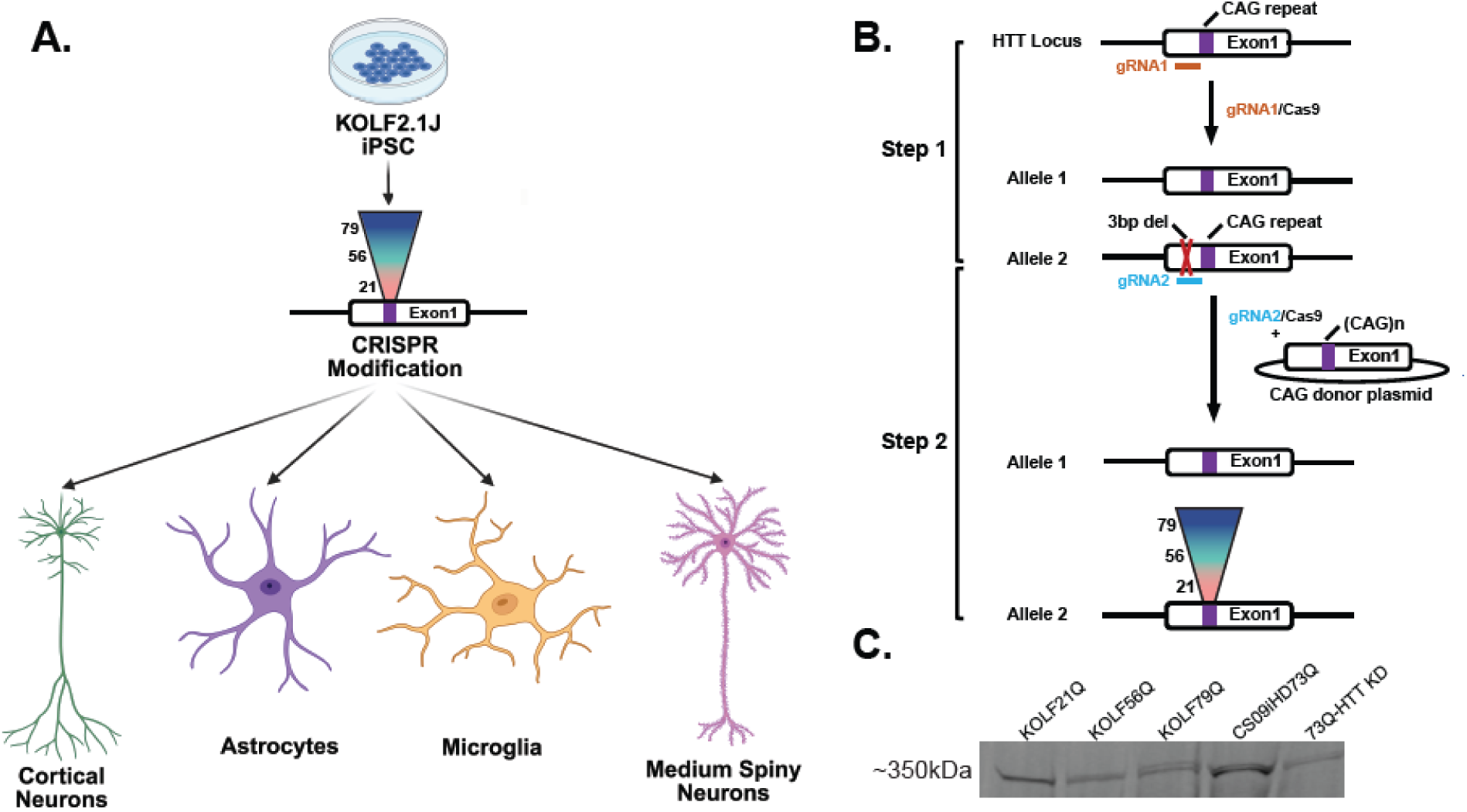
Overall Experimental Design and the Generation of the KOLF Isogenic Series. **A.** Schematic of cell types differentiated from KOLF2.1J iPSC lines after CRISPR modification. Illustration modified and created in *BioRender. Lab, T. (2026)* https://BioRender.com/d78omrg **B.** Strategy for generating the iPSC HTT CAG allelic series. CRISPR/Cas9 technology was used to introduce a heterozygous indel mutation immediately upstream of the HTT Exon1 CAG repeat region. Designing a “neo-sgRNA” to this mutated sequence allowed for allele-specific introduction of HTT Exon1 with increasing expansion of the CAG repeat region. Four clones of each genotype were evaluated by Sanger sequencing and one of each selected for full quality control and neural and glial differentiations. **C.** Expression of HTT (antibody 5526) in the KOLF iPSC lines (21Q, 56Q, 79Q) along with a positive control patient iPSC line (CS09iHD73Q) and negative control using HTT KD in the 73Q line (73Q-HTT KD) by western blot.

### NGN2 mediated cortical neurons

Given that many studies involving the iNDI lines are differentiated into NGN2 cortical-like neurons for analysis and comparisons across disease mutations, we differentiated the HD series into glutamatergic cortical neurons using the NGN2 overexpression system (Figure 2A).^20^ All three modified iPSC lines were transfected with the doxycycline inducible PB-TO-hNGN2 piggy bac^32^ plasmid and maintained under puromycin selection for plasmid expression. The cell populations were subsequently differentiated into a cortical neuron population using a protocol modified from Pantazis et. al.^18^. After 28 days, each line with 21, 56, or 79Qs expressed a high percentage of MAP2 positive mature neurons with 80-95% of nuclei NEUN positive (Figure 2B, 2C, 2D, 2E). 80% of the mature neuronal populations were also found to express the layer VI cortical neuron maker TBR1 (Figure 2C, 2E), supporting the ability of each of the isogenic lines to differentiate into NGN2-mediated cortical neurons.

**Figure 2:**
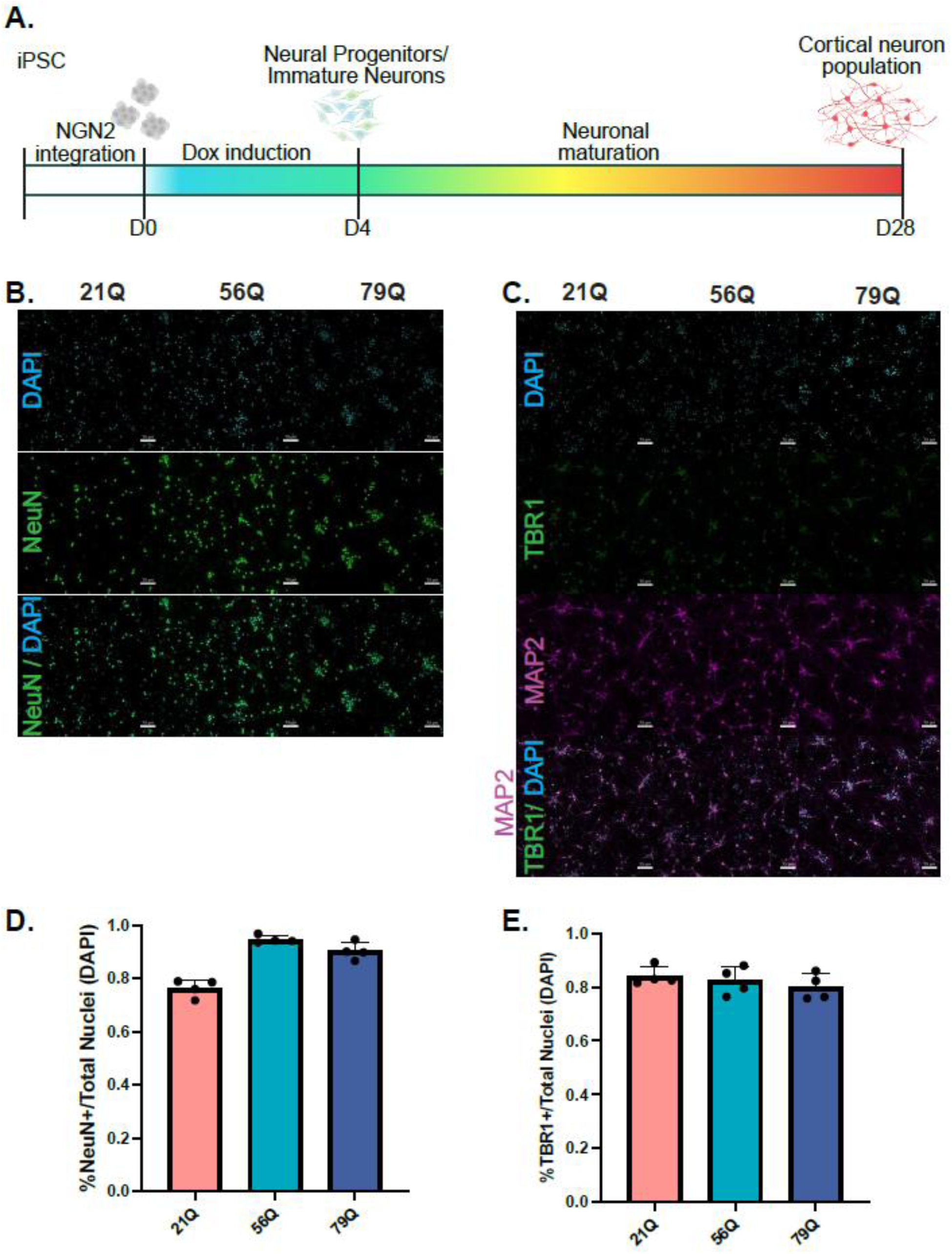
KOLF HTT Isogenic iPSCs Differentiate Into Cortical Neurons. **A.** Schematic of NGN2-cortical neuron differentiation from hiPSCs. Illustration created in *BioRender. Lab, T. (2026)* https://BioRender.com/8tkffj6. **B.** Representative images of KOLF 21Q, 56Q and 79Q NGN2 neurons with neuronal marker NeuN. Scale bar = 70 µm. **C.** Representative images of KOLF 21Q, 56Q and 79Q NGN2 neurons with neuronal markers TBR1 and MAP2. Scale bar = 70 µm. **D.** Quantification of percentage of cells expressing NeuN, normalized to DAPI. n = 4 images/line, across 1 biological replicate. **E.** Quantification of percentage of cells expressing TBR1, normalized to DAPI. n = 4 images/line, across 1 biological replicate.

### 79Q iAstrocytes display dysfunctional astrocytic properties and morphology

Astrocytes are the major glial cell types in the central nervous system (CNS) that support neurons and overall brain homeostasis. They contribute to HD through changes in the secretion of proinflammatory cytokines, increases in reactive oxygen species, withdrawal of fibers from neuronal synapses, increased extracellular potassium and decreased glutamate uptake and more.^33, 34^ The KOLF2.1J isogenic HD series were differentiated into astrocytes (iAstrocytes) using our previously published iAstrocyte differentiation protocol^22^ and express known astrocyte protein markers (GLAST, GFAP and SOX9),^35^ Figure 3A, 3B, 1 biological replicate). We previously showed that patient-derived HD iAstrocytes had dysfunctional astrocytic properties potentially arising through altered maturation, including decreased GLAST expression, aberrant morphology and decreased ability to uptake glutamate.^22^ Similarly, KOLF2.1J 56Q and 79Q iAstrocytes have decreased GLAST protein expression and a slight decrease in SOX9 protein fluorescence intensity compared to 21Q iAstrocytes (Figure 3C, 3D). HD CAG expansion iAstrocytes also have progressively decreased morphological characteristics in 56Q and 79Q compared to 21Q (Figure 3E, 3F).

**Figure 3:**
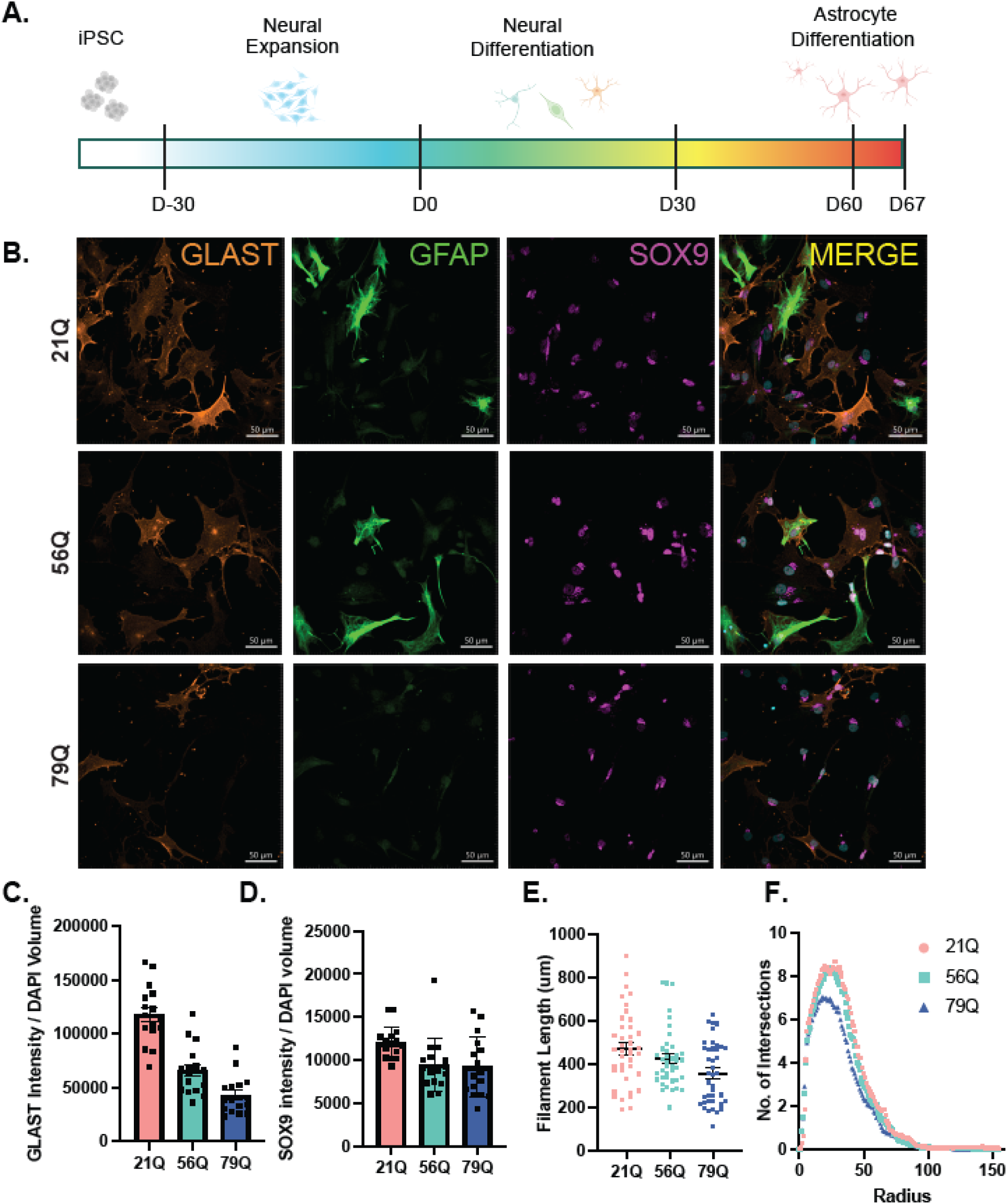
KOLF HTT Isogenic iPSCs Differentiate Into Astrocytes. **A.** Schematic of astrocyte differentiation from hiPSCs. Illustration created in *BioRender. Lab, T. (2026)* https://BioRender.com/cdldkdb. **B.** Representative images of KOLF 21Q, 56Q and 79Q iAstrocytes with astrocyte markers against GLAST, GFAP and SOX9. Scale bar = 50 µm. 1 biological replicate / line. **C.** 56Q and 79Q iAstrocytes express decreased GLAST (normalized to DAPI volume) compared to 21Q iAstrocytes. Points indicate 40x images captured. n=16 images / 40x field of view / line, 1 biological replicate / line. **D.** 56Q and 79Q have slightly decreased expression in SOX9 fluorescence intensity (normalized to DAPI volume) compared to 21Q iAstrocytes. Points indicate 40x images captured. n=16 images / 40x field of view / line, 1 biological replicate / line. **E.** 79Q iAstrocytes have decreased morphological characteristics compared to 21Q indicated by total filament length (points represent individual iAstrocytes) while 56Q have slightly decreased morphological characteristics but not as severely. n=40 iAstrocytes / line, 1 biological replicate / line. **F.** 79Q iAstrocytes have decreased number of intersections by Sholl analysis (averaged number of intersections of iAstrocytes). n=40 iAstrocytes / line, 1 biological replicate / line.

### No phagocytic changes in iMG with expanded repeats

We next differentiated the isogenic lines into the resident immune cells of the CNS --microglia–using a previously established protocol (Figure 4A).^24^ Unlike neurons and astrocytes, microglia are derived from the mesoderm rather than the neurectoderm, allowing us to examine the cell lines capacities to differentiate down another germ layer. All three repeat lengths (21Q, 56Q, and 79Q) generated microglia like cells (iMGs) expressing key microglial/macrophage proteins IBA1 and PU.1 as well as microglial surface receptors P2RY12 and CX3CR1 (Figure 4B). In addition to expressing microglial proteins and receptors, iMGs from all three repeat lengths were able to perform a basic microglial function, phagocytosis. Using a previously described assay,^24^ PH (PHrodo) sensitive fluorescently labeled S.*aureus* particles cultured with our iMGs, we examined the amount of labeled particles that entered an acidic environment (such as a phagophore) by measuring fluorescence expression over a 24-hour period. For all three lines (21Q, 56Q, and 79Q), there was a significant increase in fluorescence in our PHrodo cultures after 12, 18, and 24 hours (Figure 4C, S4) suggesting that all three lines can engulf the S.*aureus* particles and introduce them to an acidic environment for degradation as would occur in normal phagocytosis. We did not observe any significant, reproducible differences in the level of fluorescence between the different repeat lengths, consistent with previous iMG studies that have either observed no differences or inconsistent differences in the phagocytic activity of HD microglia.^16, 36^

**Figure 4:**
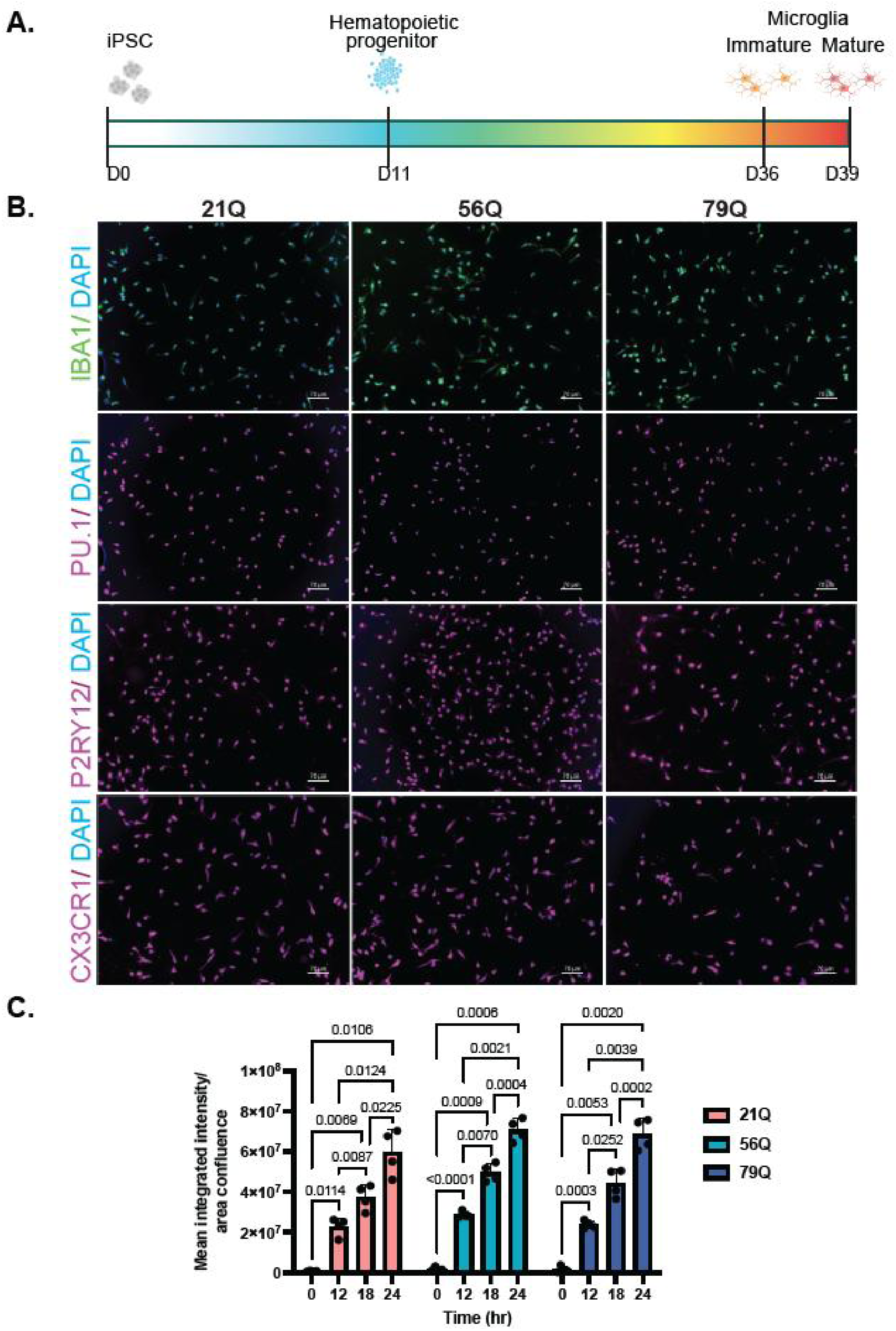
KOLF HTT Isogenic iPSCs Differentiate Into iMG. **A.** Schematic of iMG differentiation from hiPSCs. Illustration created in *BioRender. Lab, T. (2026)* https://BioRender.com/6oax5iq. **B.** Representative 20x images of 21Q, 56Q, and 79Q iMGs stained for IBA1 (green), PU.1 (magenta), P2RY12 (magenta), and CX3CR1 (magenta). All counterstained with DAPI (blue). Scale bar = 70 µM. 1 biological replicate / line. **C.** Graph representing phagocytosis levels measured by fluorescence expression (normalized to brightfield cellular confluence) at timepoints 0, 12, 18, and 24 hours in each of the 21Q, 56Q, and 79Q cell lines after being co-cultured with pHrodo labeled *S. aureus* particles. Fluorescent levels increase with time in all three lines and significant differences are observed in these levels for each line at timepoints 0, 12, 18, and 24hrs (two-way ANOVA with Bonferroni multiple comparisons correction; time factor: F(1.171,10.54)= 660.8, P<0.0001, individual p values indicated on graph for significant comparisons). No significant differences were observed in the fluorescent levels between the cell lines for any given time point (two-way ANOVA with Bonferroni multiple comparisons correction; line factor: F(2, 9)= 2.984, P=0.1014). Data shown represents 1 of 3 replicate experiments for a subset of timepoints additional experimental data can be found in Figure S4), Data points = 4 images averaged per well from n = 4 wells per cell line.

### HTT isogenic and KO lines can differentiate into neurons enriched for striatal neurons

We next generated homozygous and heterozygous HTT knockout lines to provide an isogenic system in the KOLF2.1J background in which to study loss of HTT normal function compared to expanded repeat lines, similar to studies carried out previously in the RUES2 isogenic lines.^12, 15^ To generate HTT KO lines, the HTT locus was targeted for deletion of the entire coding sequence (Figure 5A). This approach produced only heterozygous knockouts; therefore, the pool of heterozygous knockouts was re-targeted for deletion of the second allele. Three homozygous KO clones were identified by PCR and Sanger sequencing, and two were chosen for Western blot to confirm lack of protein expression (Figure 5B, S5A-C); both *HTT*KO_1, *HTT*KO_2 showed absence of HTT protein. Both clones were further confirmed pluripotent by qPCR for standard markers (Figure S5D) and karyotypically normal by aCGH analysis (Figure S6).

**Figure 5:**
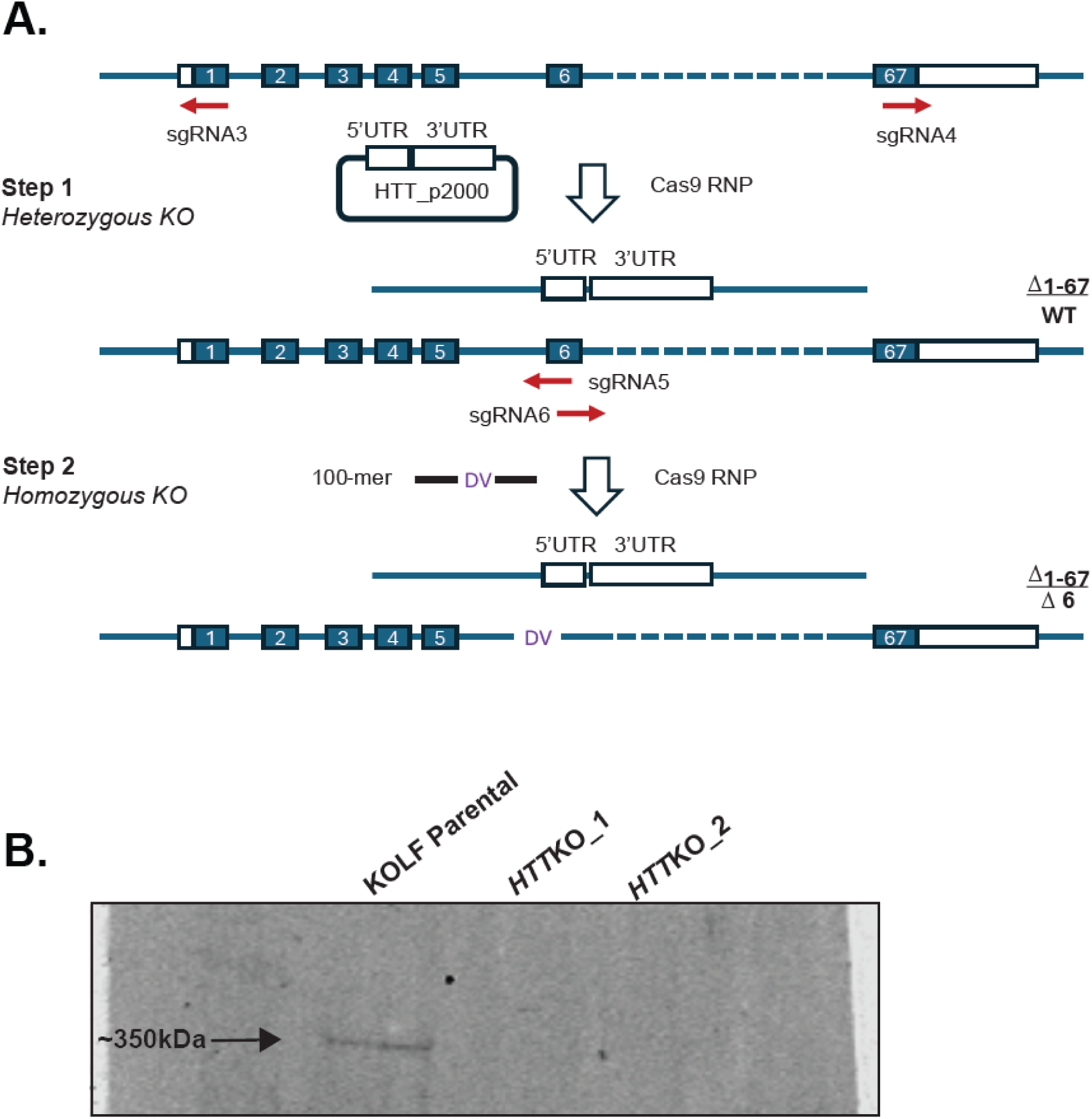
Generation of KOLF HTT Knockout Lines. **A.** Schematic of the generation of the KOLF HTT KO lines and steps from heterozygous KO (Step 1) to homozygous KO (Step 2). **B.** HTT is successfully knocked out from the KOLF parental iPSC line determined by western blot analysis (antibody 5526).

Medium spiny neurons (MSNs) are the most overtly affected cell type in human HD. The KOLF iPSC lines (21Q, 56Q, 79Q, *HTT*KO_1, *HTT*KO_2) were differentiated for 37 days into mature neurons enriched for MSNs as described (Figure 6A).^25^ At Day 37, all lines produced a high percentage of mature neurons as evidenced by 70-90% expression of the mature neuron marker NeuN (Figure 6B, 6C, S7A), and high MAP2 expression (Figure S7B). They also displayed high expression of FOXP1, a transcription factor important for striatal neuron development. There does not appear to be any difference in their ability to mature as there were no significant differences in NeuN expression levels between lines (Figure 6C). Here we defined MSNs by co-expression of DARPP-32 and CTIP2 (alternatively known as BCL11B); all lines produced 35-60% DARPP-32+/CTIP2+ cells as expected.^25^ There was some variance in the levels of DARPP-32+/CTIP2+ co-expression between the lines, with the expanded lines having slightly increased expression (56Q and 79Q) and the *HTT*KO lines having slightly decreased expression, which was significantly different when *HTT*KO lines were compared to the expanded repeat lines but not when compared to control (21Q) (Figure 6D). We next asked if the reduction in the number of observed MSN’s in the *HTT*KO lines influenced their potential to form synapses by measuring the co-localization of presynaptic protein SYP and post-synaptic protein PSD95 puncta as a measurement of synapse number. There were no significant differences, although the *HTT*KO_2 appears to have lower co-localization of SYP and PSD95 compared to 21Q which will need to be further investigated (Figure S8A-B).

**Figure 6:**
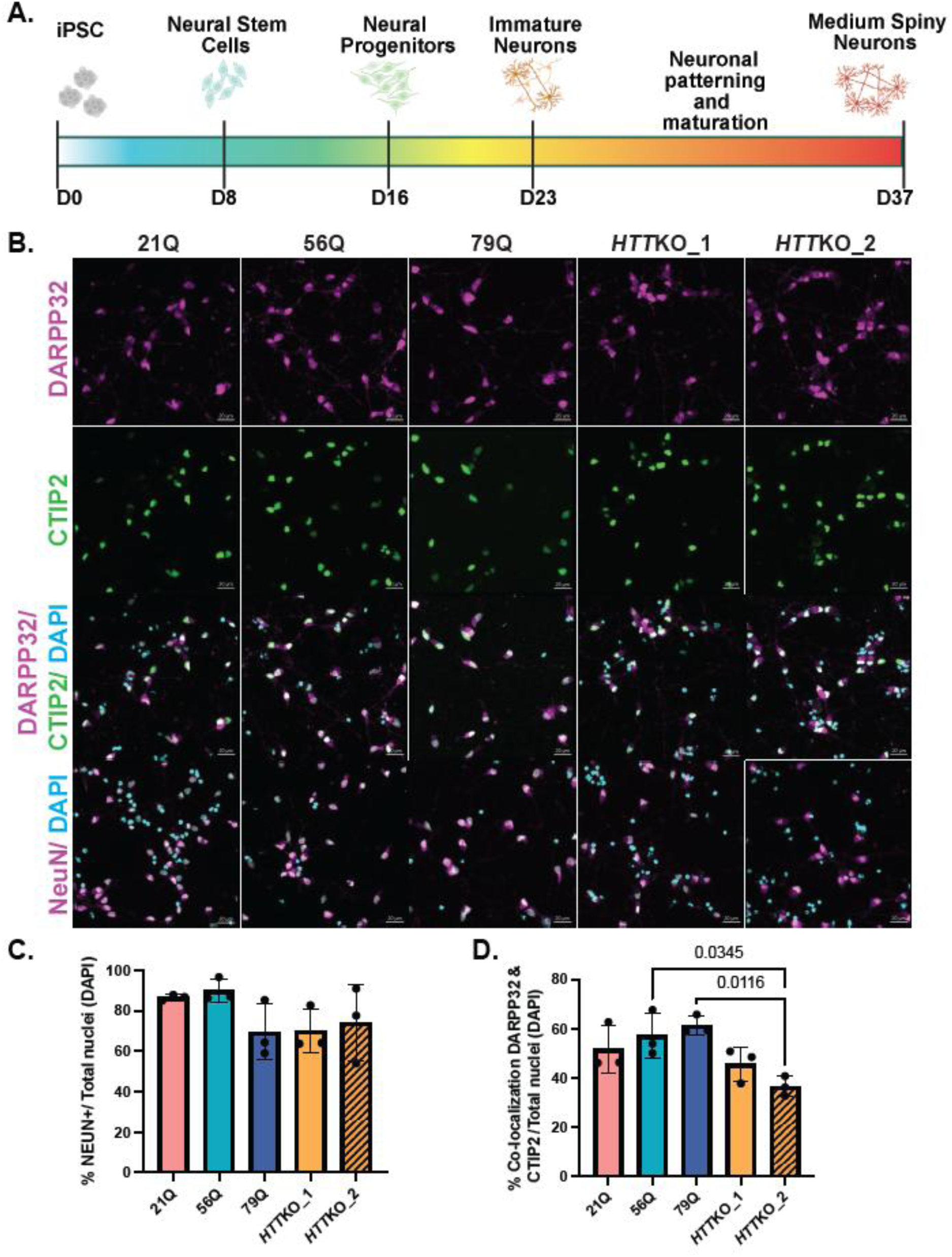
KOLF HTT Isogenic iPSCs Differentiate Into Striatal Neurons. **A.** Schematic of striatal neuron differentiation from hiPSCs. Illustration created in *BioRender. Lab, T. (2026)* https://BioRender.com/0k5xuho. **B.** Representative 20x images of KOLF 21Q, 56Q, 79Q, *HTT*KO_1, and *HTT*KO_2 with mature neuron marker NeuN, and striatal neuron markers DARPP-32 and CTIP2. Scale bar = 20 µm. **C.** Quantification of NeuN, normalized to DAPI. n = 3 images/line, across 3 biological replicates. **D.** Quantification of DARPP-32/CTIP2 colocalization, normalized to DAPI. n = 3 images/line, across 3 biological replicates.

### HTT KO lines display similar deficits as long CAG repeats

To establish baseline transcriptomes and evaluate the effects of mHTT and HTT KO on the coding transcriptome, we performed mRNA-seq on the MSNs generated from our lines. Unbiased clustering analysis, or principal Component Analysis (PCA), was used to explore clustering based on gene expression. We observed that replicates for each cell line clustered together and also clustered by genotype with both CAG expansion lines (56Q and 79Q) and HTT KO lines (*HTT*KO_1 and *HTT*KO_2) clustering separately from control lines (21Q) (Figure 7A). Differential gene analysis was performed and identified 96 differentially expressed genes (DEGs) for the 56Q vs 21Q, 389 DEGs in 79Q vs 21Q, and 1441 DEGs in *HTT*KO vs 21Q comparisons (Figure 7B, S9A-C, Table S4). There was minimal overlap in DEGs between comparisons (Figure 7B). Therefore, ingenuity pathway analysis (IPA) was used to investigate biological changes at the pathway level on identified DEGs (Figure 7C). Many of the observed pathways are dysregulated in HD systems including CREB Signaling in Neurons^37, 38^, G-Protein Coupled Receptor Signaling^39^ and Focal Adhesion Kinase (FAK) Signaling^40^ (Figure 7C, S9D-F). From this analysis, we observed that the DEGs in the 79Q and KO striatal neurons compared to controls are more similar than the 56Q line compared to controls at the pathway level. (Figure S9D-F). We have previously observed similarities between expanded repeats and KO alleles^15^ and distinct profiles for highly expanded repeats and adult-onset repeats^25^. Surprisingly however, the 56Q line showed dysregulation in similar pathways to the 79Q and HTTKO lines, but in the opposite direction in most cases (Figure 7C).

**Figure 7:**
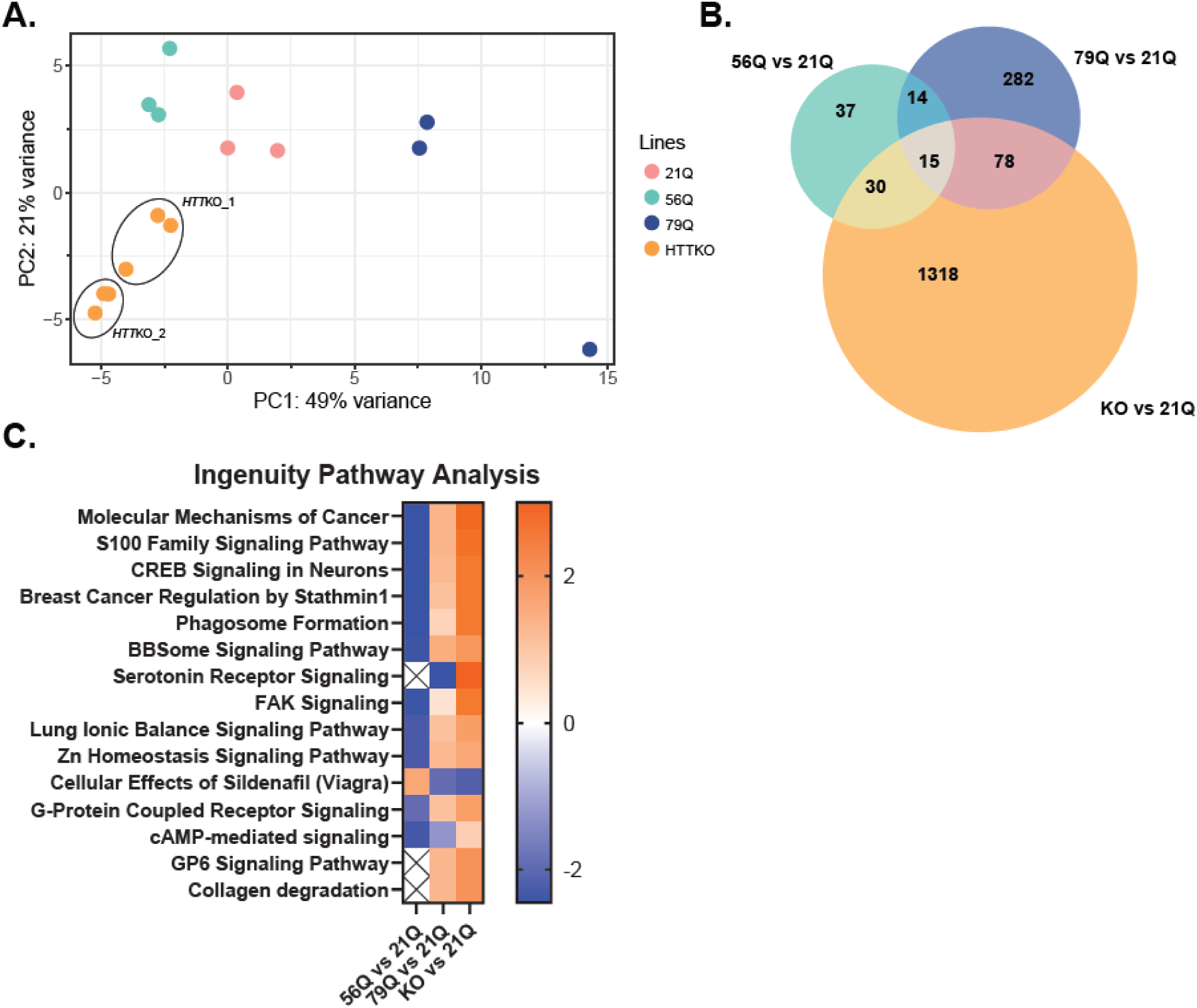
mRNAseq analysis of KOLF Isogenic Series and KO Lines Striatal Neurons. **A.** Principal component analysis (PCA) of rlog normalized counts from differentiated striatal neurons with clear separation between the lines (21Q, 56Q, 79Q, *HTT*KO_1, and *HTT*KO_2) and minimal variability between line replicates. **B.** Number of differentially expressed genes (DEGs) and comparisons between 21Q, 56Q and 79Q (adjusted p value < 0.05). **C.** Ingenuity Pathway Analysis (IPA) of DEGs showing top categories and activation scores for 56Q vs 21Q, 79Q vs 21Q and KO vs 21Q with 79Q comparison similar to KO pathways.

## Discussion

Here, we generated, differentiated, and characterized in selected assays a set of isogenic iPSC lines for HD in the KOLF2.1J background, providing a resource for the community that can be used for both the study of HD mechanisms as well as directly compare to other neurodegenerative disease mutations available through the iNDI collection. The lines can successfully differentiate into neurons (cortical and striatal) and glia (astrocytes and microglia), and we identified transcriptomic and phenotypic alterations consistent with those observed in other HD patient-derived lines in selected assays. Based on protein expression, all lines differentiated into the intended cell types. In selected assays, there was no difference in basic iMG function, however, iAstrocytes and neurons enriched for MSNs displayed disrupted phenotypes.

79Q iAstrocytes exhibited dysfunctional GLAST expression and altered morphology, mirroring abnormalities previously observed in HD patient-derived iAstrocytes. These changes may reflect impaired glutamate uptake or an inability to achieve full astrocyte maturation, consistent with reports that mHTT disrupts development and cellular differentiation.^14, 22, 41, 42^ A more comprehensive characterization—including protein markers such as GFAP, S100B, NFIA, and FABP7, along with assays assessing inflammatory responses, developmental state, and potassium-buffering capacity will further inform mechanisms. Notably, the 79Q iAstrocytes showed progressively greater dysfunction than the 56Q line as expected. Future studies will evaluate whether HD iAstros can provide synaptic support to neurons and whether neurons can rescue iAstro phentoypes in co-culture studies.

Neither 56Q nor 79Q iMG displayed impairment in protein marker expression or phagocytic capabilities. HD Microglia have been observed to be more reactive or inflammatory in HD patient brains^43, 44^ and mouse models^45, 46^ and the expression of mHTT alone was found to produce a pro-inflammatory transcriptome in an immortalized microglia model.^46^ Consequently, an effect on microglial immune cell function might be expected; however, we did not detect any consistent differences in one such function, phagocytosis of a pathogen (S. *aureus*). This is consistent with previous data for stem cell derived HD microglia where either no differences were observed between HD and control cells or observed differences seemed to be specific to individual cell lines and may not reflect an HD effect but rather cell line variability.^16, 36^ The effects of the HD mutation on microglia may be more subtle or relate more to their interactions with and maintenance of other cell types such as neurons and astrocytes. Further research using more complex cell models, combining microglia with other cell types, could help define any potential effects mHTT expression has on HD microglia and how they may contribute to disease progression.

Because degeneration of MSNs in the basal ganglia is a major hallmark of HD,^47^ we differentiated the isogenic KOLF lines into neuronal cultures enriched for MSNs and conducted both staining-based characterization and transcriptomic analysis. To distinguish HTT loss-of-function effects from mHTT gain-of-function effects, we also generated homozygous HTT knockout iPSC lines, with additional characterization still required for the heterozygous HTT KO line. All lines readily differentiated as indicated by a high percentage of DARPP-32 and CTIP2 co-localization (30-60%)^25^ and no detectable changes in NeuN or synapse number. Among all lines, one KO line, *HTT*KO_2, showed reduced DARPP-32/CTIP2 co-localization, further supporting the essential role of HTT in neuronal development.^41, 48^ Downstream implications of mHTT gain or HTT loss include an impact on early embryo patterning, ciliogenesis, synaptic density, signaling pathways and cell-cell communications that can be investigated using the KOLF lines.^14, 41^

mRNA-seq gene expression analysis of 56Q, 79Q, and HTT KO striatal neurons relative to 21Q revealed that although these lines affect many of the same pathways, they do so through distinct gene expression patterns and in some instances, in alternating regulatory directions. Our data highlights dysregulated pathways related to CREB signaling, cell adhesion, ion homeostasis, G-protein–coupled receptor signaling, and cAMP-mediated signaling. The sets of DEGs contributing to these pathways differed across the lines, indicating that multiple mechanisms of dysregulation operate depending on CAG repeat length and potentially on whether the perturbation reflects HTT loss or mHTT gain of functions. Transcriptionally, our 79Q striatal neurons more closely resemble the KO striatal neurons than either the 21Q neurons or 56Q neurons and show concordant overactivation of several pathways. This overlap in transcriptional dysregulation observed between CAG expansion and HTT KO was previously observed in iPSC derived cortical neurons using the RUES2 isogenic cell series^15^, highlighting how some dysregulation seen with CAG expansion may be a consequence of HTT loss of function. Surprisingly, the transcriptional dysregulation observed in our 56Q striatal neurons were not as concordant as those between the 79Q and *HTT*_KO lines and while similar pathways were affected, these pathways were primarily downregulated in the 56Q neurons. Here, the 56Q line may be representative of an earlier point of pathway dysregulation. We speculate that the highly expanded 79Q produces a more severe HTT loss of function that recapitulates aspects of the HTT KO phenotype whereas the 56Q line shows a moderate transcriptional effect within the iPSC-derived neurons, which do not carry the chronic effects of stress or aging.

The HD KOLF2.1J isogenic series serves as a community resource, providing an isogenic series to address mechanistic questions in HD. Additionally, this series can be compared with other genetically based neurodegenerative disease models generated through the iNDI Consortium, including those for Alzheimer’s disease, Parkinson’s disease, and amyotrophic lateral sclerosis (ALS).

## Acknowledgments/Funding

This project was funded by NIH iNDI program (ZIA AG000535) and the Chan Zuckerberg Initiative together with the following NIH grants: R35NS116872 (L.M.T.), R01NS089076 (L.M.T.) and F31NS134306-01A1 (M.S.B.). Additional support was provided by HD-CARE, CIRM EDUC4-12822 (M.S.B. and N.R.M.), ARCS Foundation Orange County (M.S.B), NINDS/NIH T32 (NS082174 & NS121727-01, J.T.S.) and The Rose Hills Foundation (J.T.S.). This work was possible, in part, through access to the UCI Genomics Research and Technology Hub (GRT) of the Cancer Center Support Grant (P30CA62203). This research was supported, in part, through the Center for Alzheimer’s Disease and Related Dementias (CARD) at the National Institutes of Aging (NIA) at the NIH.

## Author contributions

Conceptualization: L.S., L.M.T., W.C.S; Data curation: L.S., J.S., M.S.B, K.Q.W.; Data analysis: R.M., M.S.B., K.Q.W., J.T.S., L.S., L.H; Funding acquisition: L.M.T; Investigation: L.S., J.T.S., K.Q.W., M.S.B., G.C., L.H.; Methodology: A.R.K., L.S., W.C.S, J.T.S., M.S.B., K.Q.W., G.C. Project administration: L.S., B.S., M.W.; Resources M.W.; Software: R.M.; Supervision: L.M.T., W.C.S.; Validation: L.H., N.R.M.; Visualization: L.S., J.S., K.Q.W., M.S.B., R.M.; Writing – original draft: L.S., J.S., K.Q.W., M.S.B., L.M.T; Writing – review & editing: all authors

## Statements and Declarations

### Statement of ethics approval

There are no human participants in this article and informed consent is not required.

## Data Availability

All data is included in the manuscript. RNAseq data will be deposited in GEO (<u>Gene Expression Omnibus</u>).

## SUPPLEMENTAL INFORMATION

**Figure S1:**
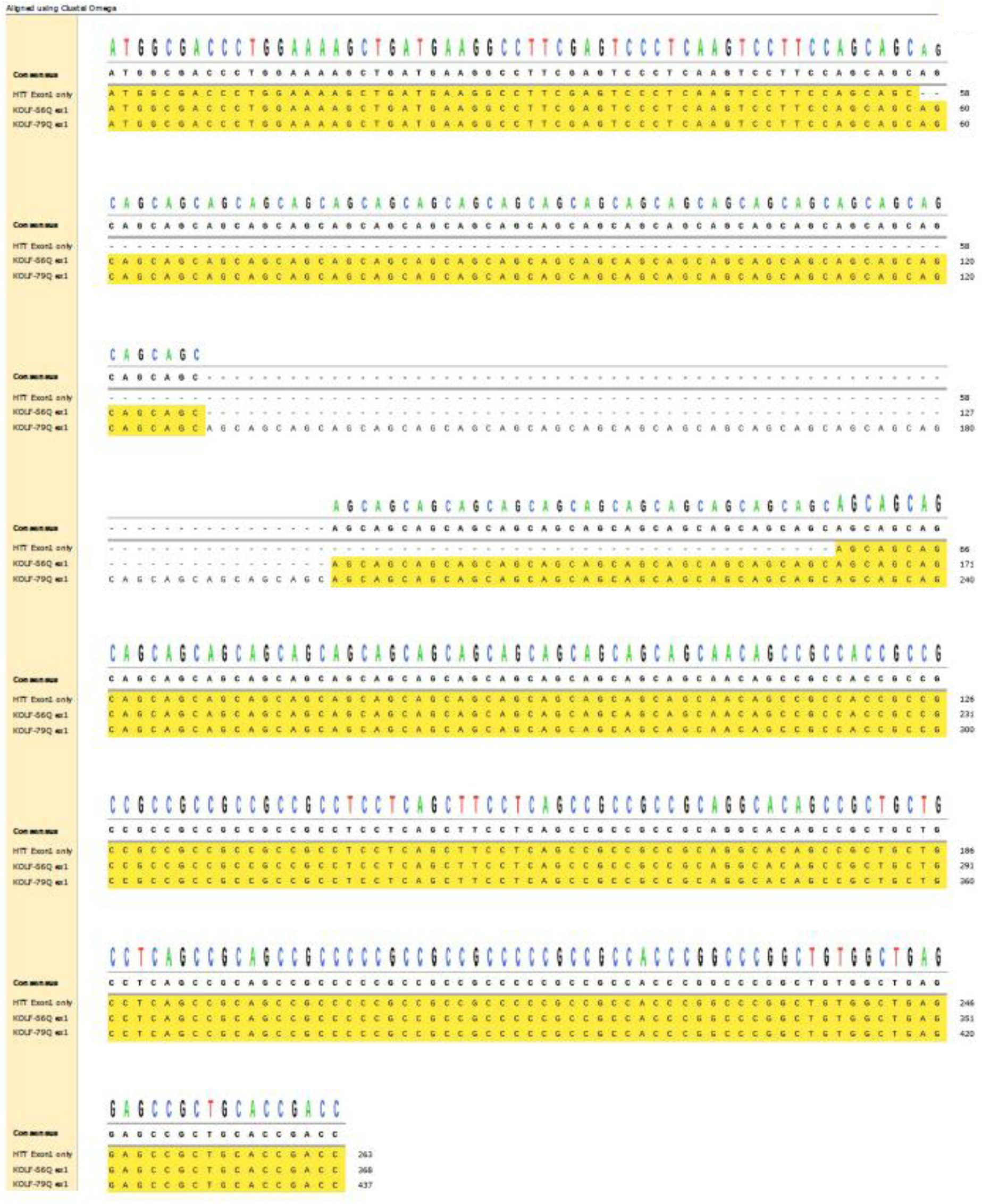
Sequencing of KOLF iPSCs. Expanded alleles (56Q and 79Q) amplified and sequenced to confirm repeat length.

**Figure S2:**
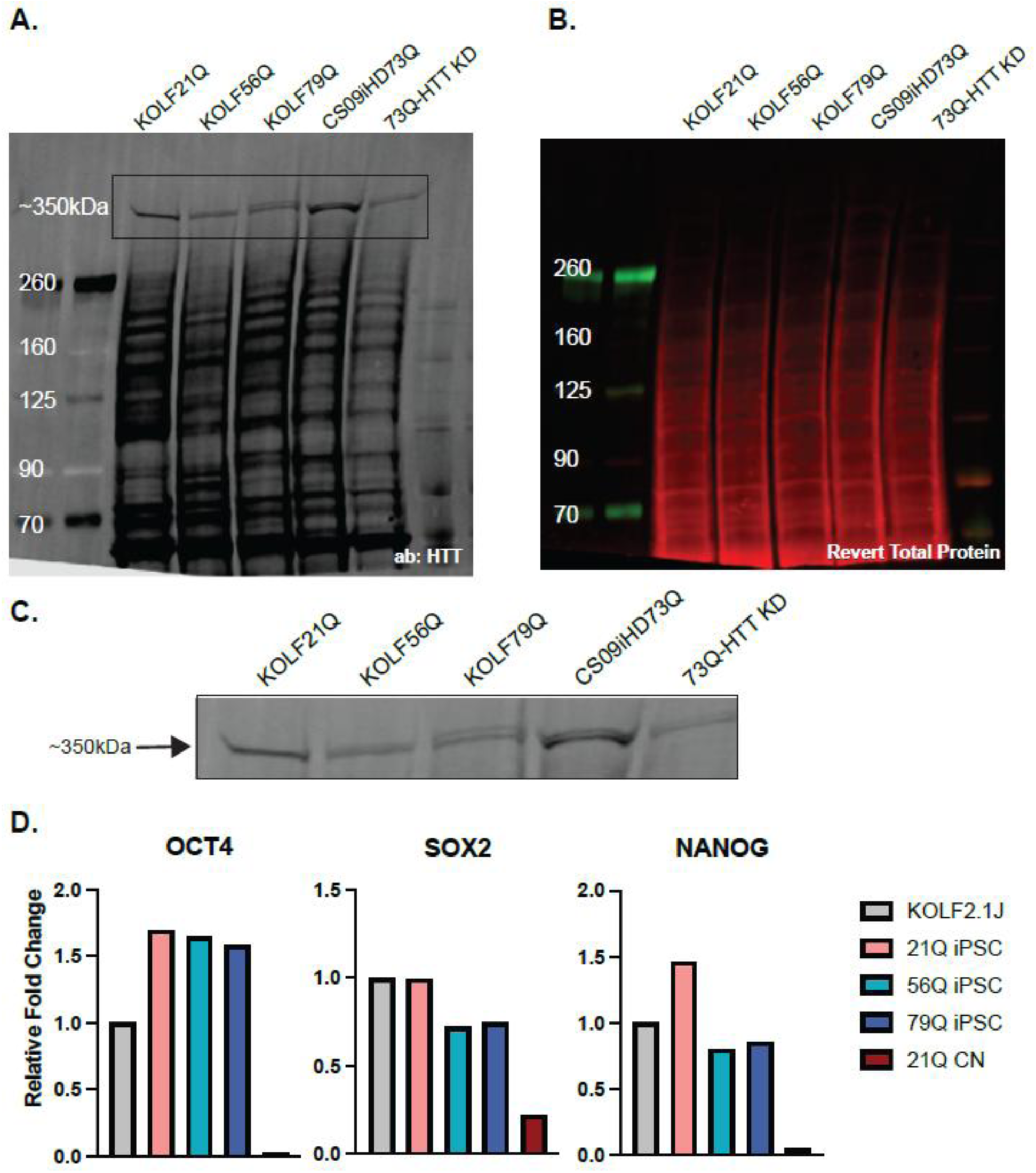
Quality control for KOLF iPSCs. **A.** Full western blot image of HTT (antibody 5526) in the KOLF iPSC lines (21Q, 56Q, 79) along with a positive control patient iPSC line (CS09iHD73Q) and negative control using HTT KD in the 73Q line (73Q-HTT KD). **B.** Revert total protein of western blot for normalization. **C.** Enlarged image of western blot used in Figure 1C. **D.** KOLF iPSCs express pluripotency markers by qPCR.

**Figure S3:**
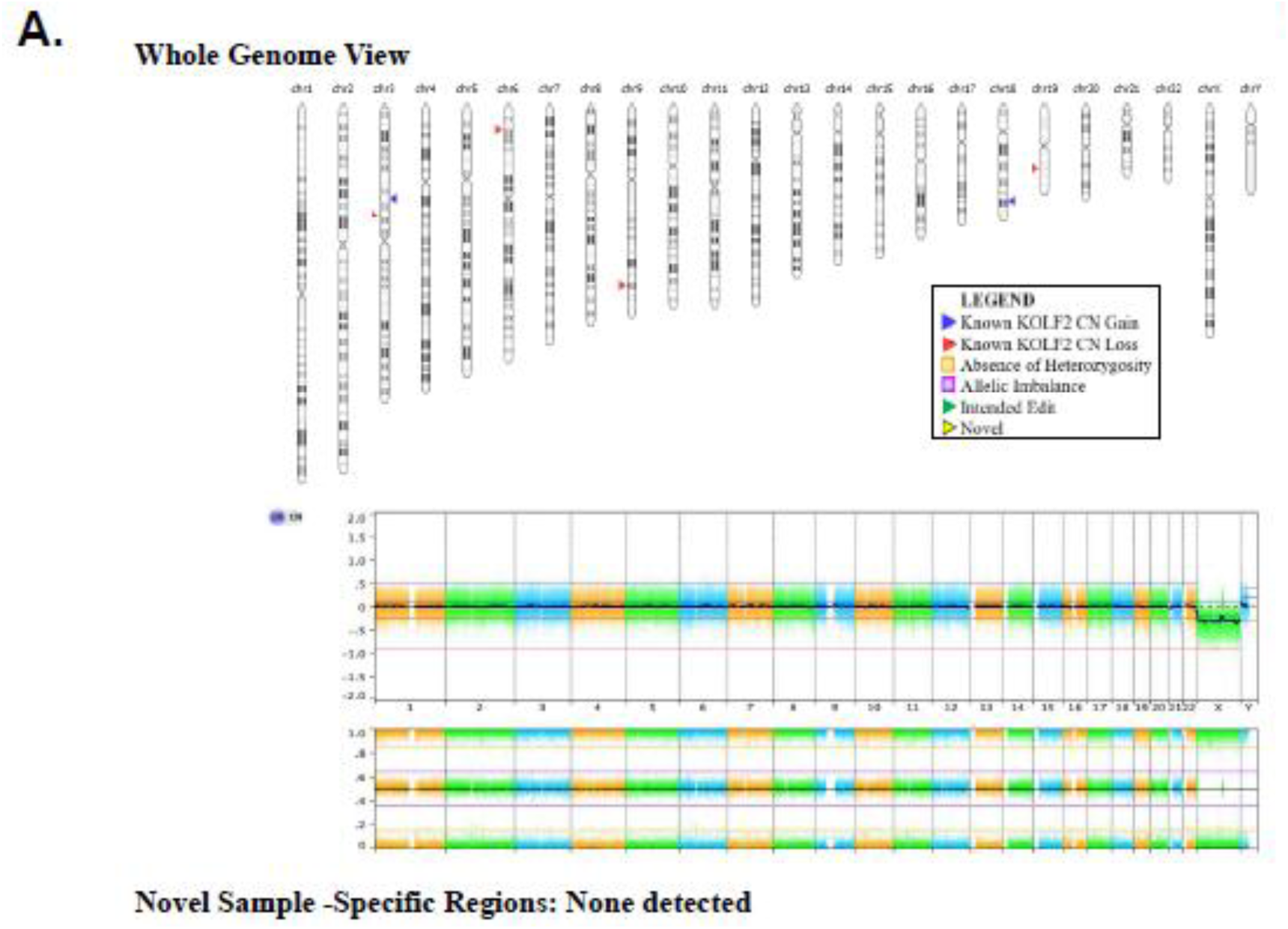
SNP Array for KOLF Isogenic Series. SNP analysis confirmed known CNVs present in the KOLF2.1J line and the original KOLF2-C1 iPSC line from which it was generated. No novel CNVs were detected. Representative result, all lines (21Q, 56Q, and 79Q) yielded the same result. SNP Analysis.

**Figure S4:**
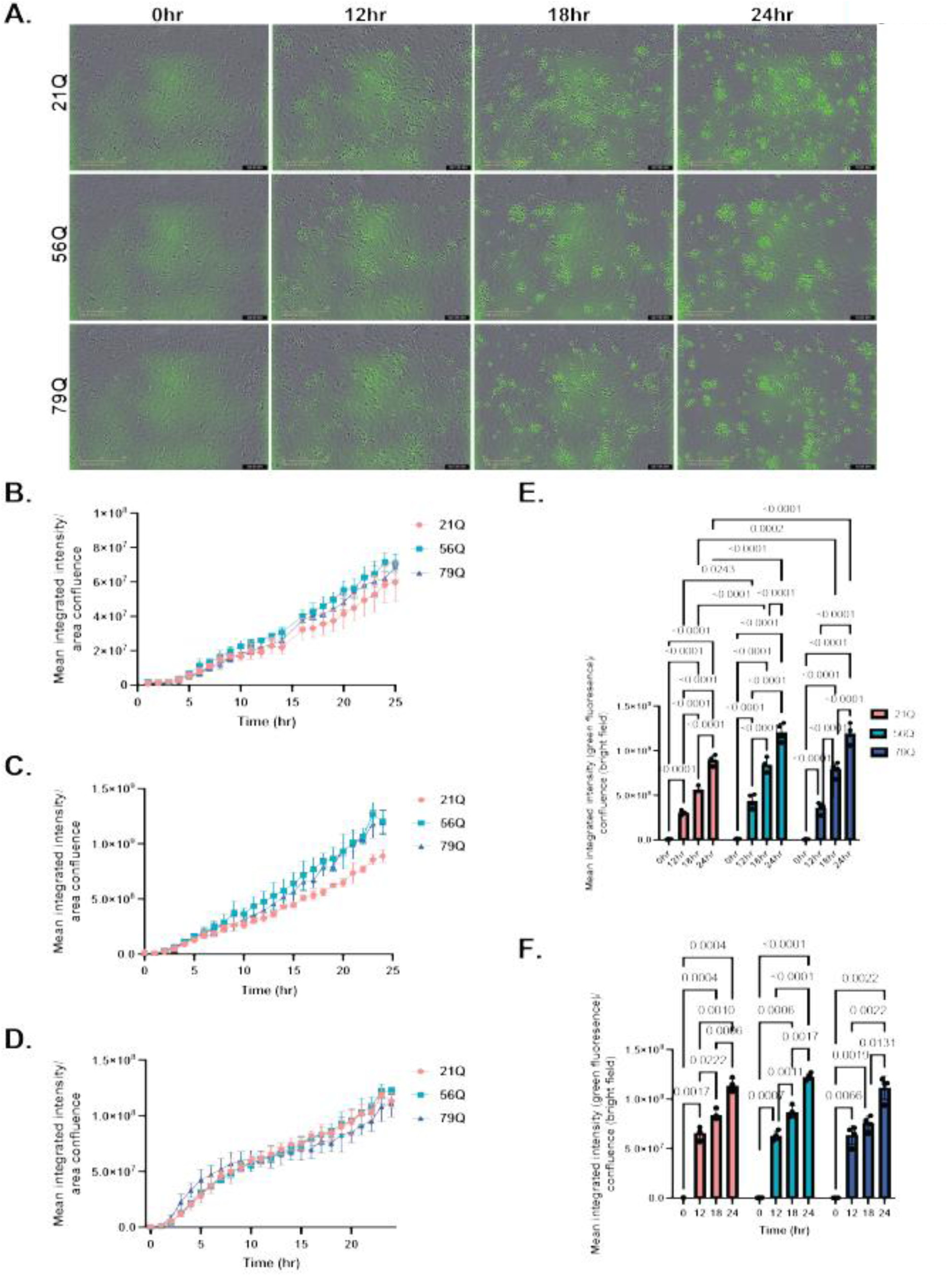
iMG PHrodo assay for phagocytosis. **A.** Representative 20x images of fluorescence levels observed the 21Q, 56Q, and 79Q lines at time points 0, 12, 18, and 24 hours for the phagocytosis assay shown in Figure 4. Scale bar = 200uM. **B.** Graph representing fluorescence levels (normalized to brightfield cell confluence) for each line at hour across 24 hrs for the phagocytosis experiment shown in Figure 4. Error bars represent the mean +/- S.D. (16 hour timepoint images were not properly focused and were removed for all lines) **C.** Graph representing fluorescence levels (normalized to brightfield cell confluence) for each line at hour across 24 hours for a second replicate phagocytosis experiment (replicate experiments represent separate rounds of parallel differentiations for all lines) . Error bars represent the mean +/- S.D. **D.** Graph representing fluorescence levels (normalized to brightfield cell confluence) for each line at hour across 24 hours a second replicate experiment (replicate experiments represent separate rounds of parallel differentiations for all lines) . Error bars represent the mean +/- S.D. **E.** Graph representing fluorescence expression (normalized to brightfield cellular confluence) at timepoints 0, 12, 18, and 24 hours each line in the phagocytosis experiment shown in panel C. Fluorescent levels increase with time in all three lines and significant differences are observed in these levels for each line at timepoints 0, 12, 18, and 24hrs (two-way ANOVA with Bonferroni multiple comparisons correction; time factor: F(3, 12)= 644.1, P<0.0001, individual p values indicated on graph for significant comparisons). Significant differences were observed in the fluorescent levels between the expanded CAG repeat lines and the 21Q lines, with the 56Q and 79Q lines showing increased fluorescence over the 21Q line for timepoints 12, 18, and 24 hours (two-way ANOVA with Bonferroni multiple comparisons correction; line factor: F(2, 24)= 33, P<0.0001). This difference between cell lines was not able to be replicated in either of the other replicate experiments. Data points = 4 images averaged per well from n=4 wells per cell line. **F.** Graph representing fluorescence expression (normalized to brightfield cellular confluence) at timepoints 0, 12, 18, and 24 hours each line in the phagocytosis experiment shown in panel C. Fluorescent levels increase with time in all three lines and significant differences are observed in these levels for each line at timepoints 0, 12, 18, and 24hrs (two-way ANOVA with Bonferroni multiple comparisons correction; time factor: F(1.753, 15.78)= 1522, P<0.0001, individual p values indicated on graph for significant comparisons). No significant differences were observed in the fluorescent levels between the cell lines at each of the 12, 18, and 24 hour time points (two-way ANOVA with Bonferroni multiple comparisons correction; line factor: F(2, 9)= 1.083, P=0.3791). Data points = 4 images averaged per well from n=4 wells per cell line.

**Figure S5:**
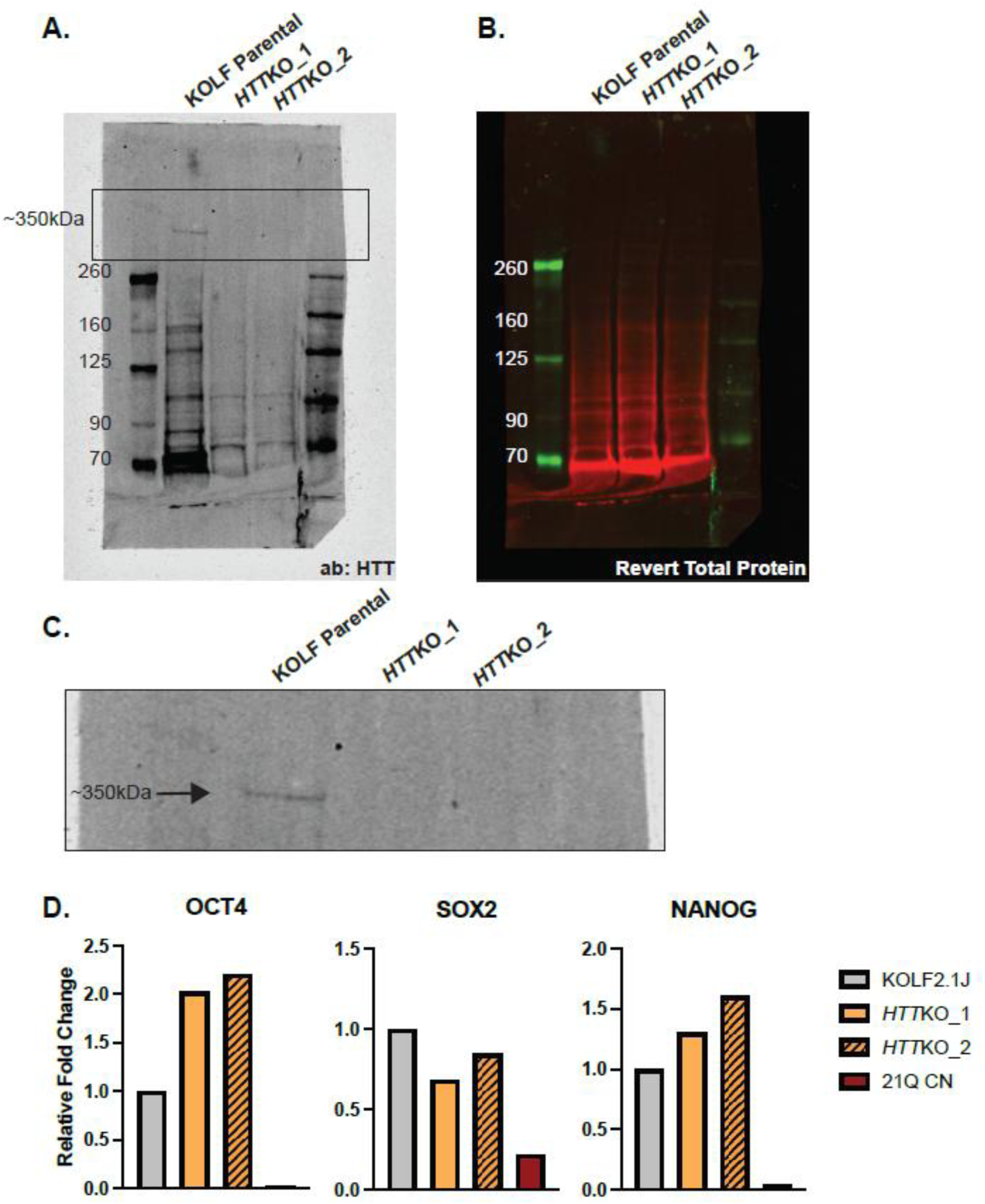
Quality control for KOLF *HTT* KO lines. **A.** Full western blot image of HTT (antibody 5526) in the KOLF KO lines compared to the parental line. **B.** Revert total protein of western blot for normalization. **C.** Enlarged image of western blot used in Figure 5B. **D.** KOLF *HTT* KO iPSCs express pluripotency markers by qPCR.

**Figure S6:**
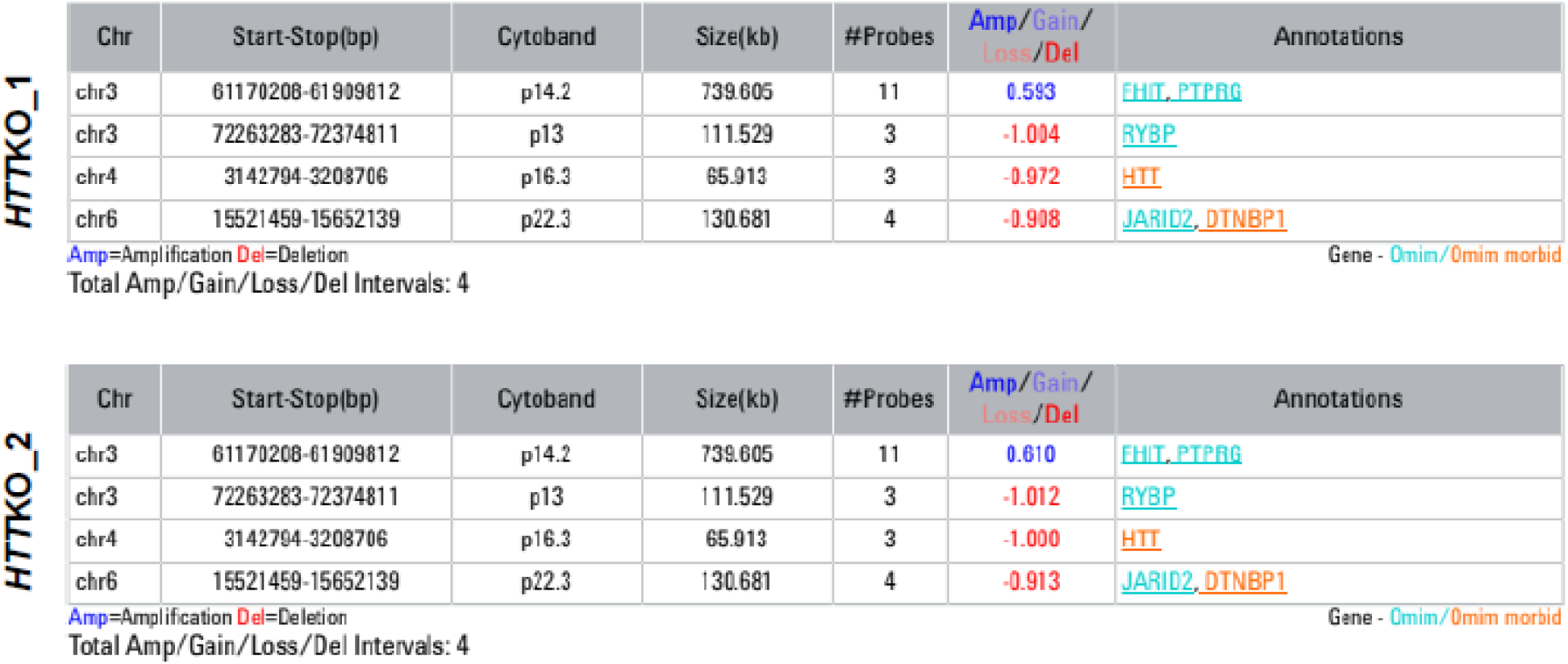
aCGH for KOLF *HTT* KO iPSCs. Normal karyotyping of *HTT*KO_1 and *HTT*KO_2 lines determined with Array Comparative Genomic Hybridization (aCGH).

**Figure S7:**
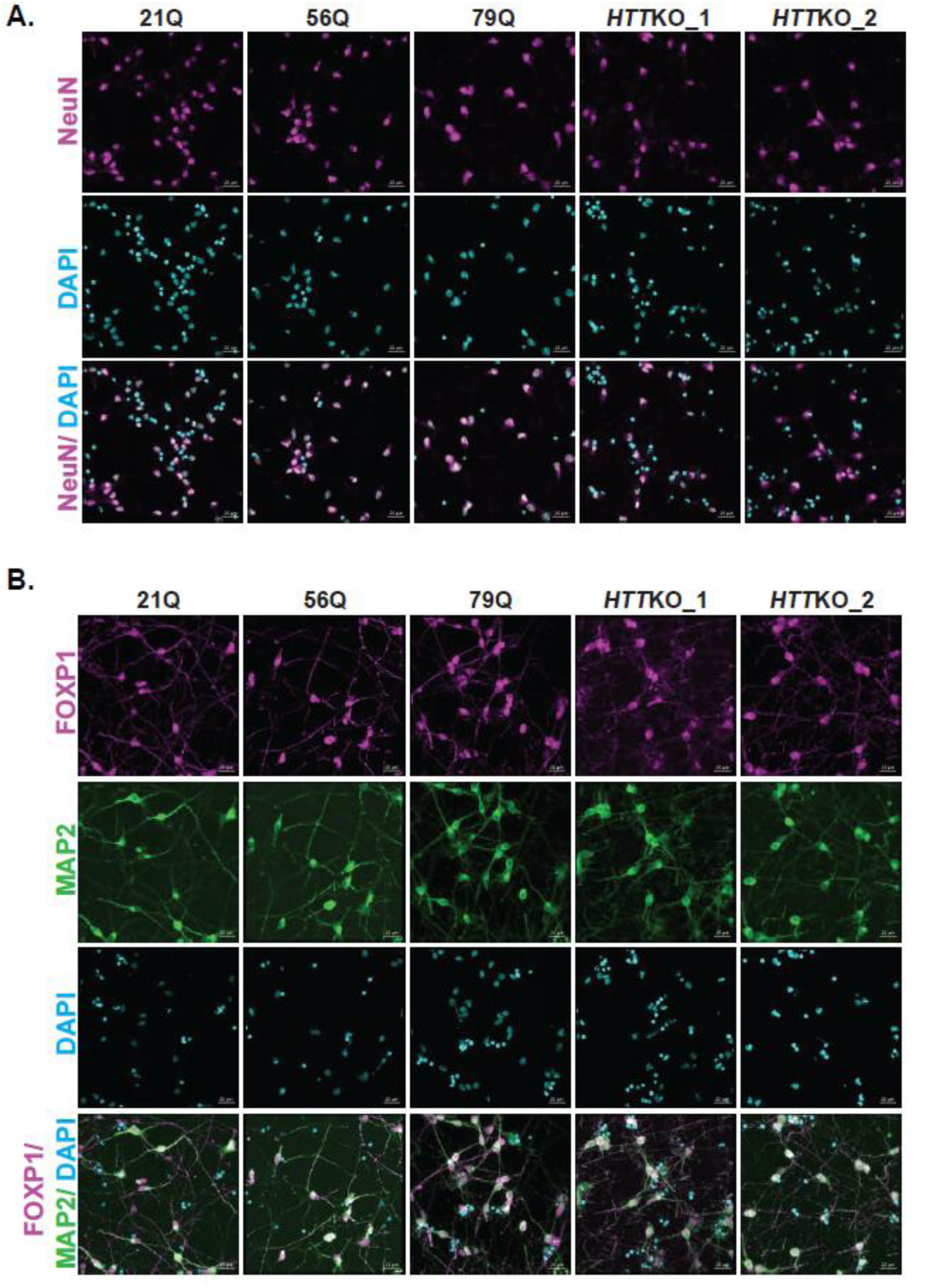
Staining of mature neurons enriched for MSNs. **A.** Representative 20x images of KOLF 21Q, 56Q, 79Q, *HTT*KO_1, and *HTT*KO_2 showing expression of MAP2 and FOXP1. Scale bar = 20 µm. **B.** Individual channels of NEUN/DAPI of KOLF 21Q, 56Q, 79Q, *HTT*KO_1, and *HTT*KO_2 shown in Figure 6B. Scale bar = 20 µm.

**Figure S8:**
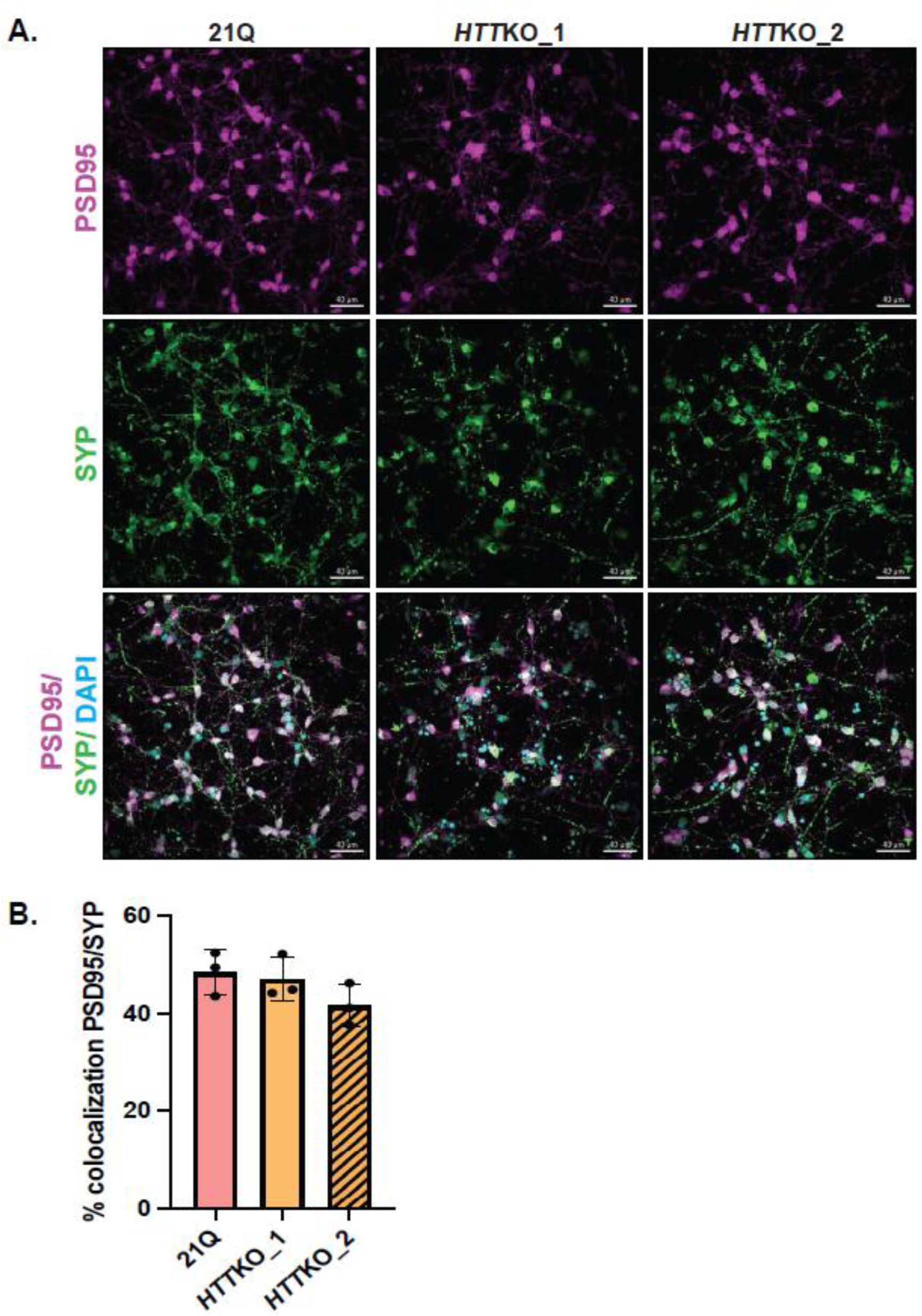
Staining of mature neurons enriched for MSNs. **A.** Representative 40x images of KOLF 21Q, 56Q, 79Q, *HTT*KO_1, and *HTT*KO_2 showing expression of pre- and post- synaptic markers SYP and PSD95, respectively. Scale bar = 40 µm **B.** Quantification of PSD95/SYN colocalization in KOLF 21Q, *HTT*KO_1, and *HTT*KO_2. n = 4 images/line, across 3 biological replicates.

**Figure S9:**
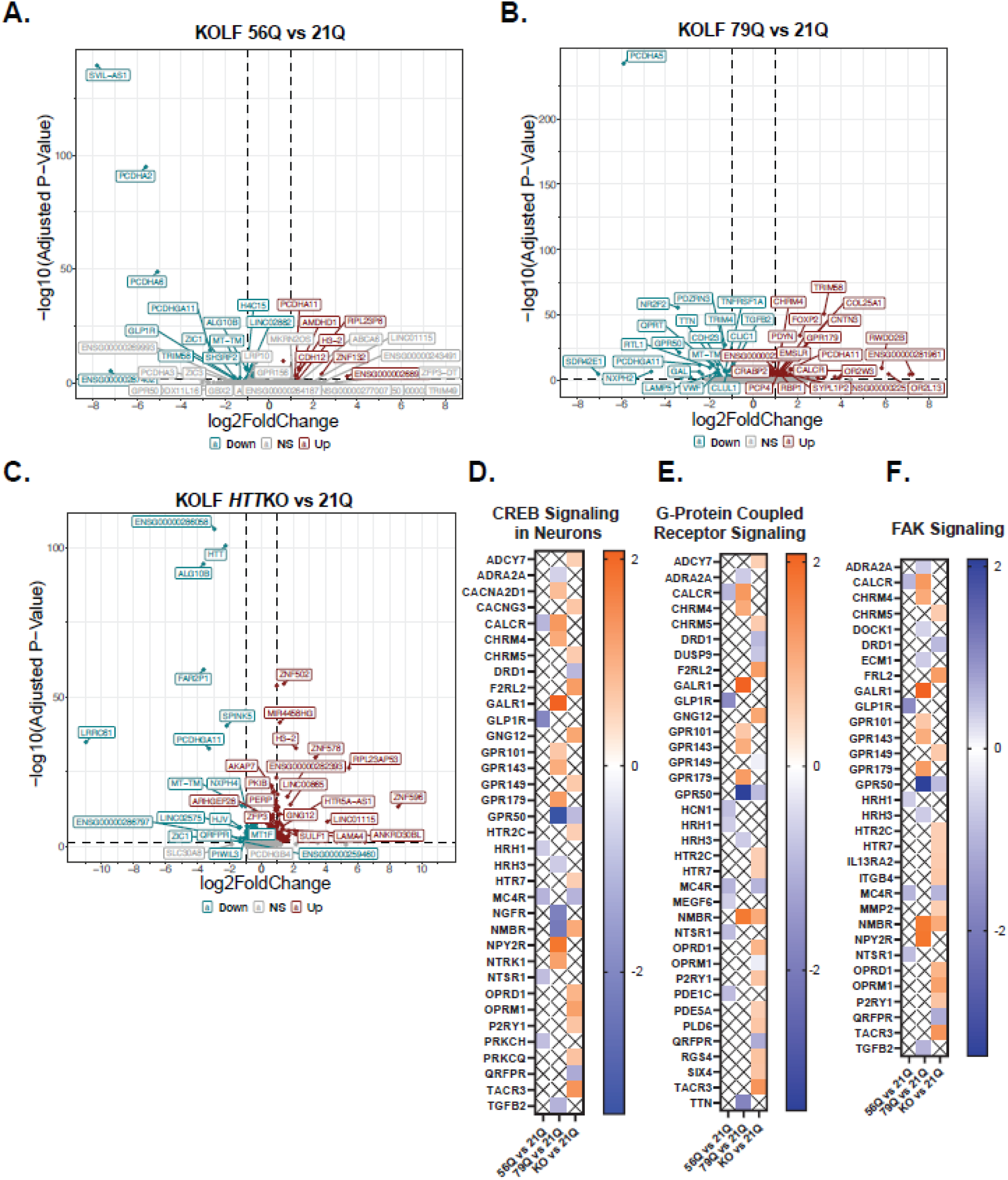
Pathway DEGs linked to Figure 7 (ie – CREB signaling, GPCR signaling, FAK signaling) **A.** Volcano plot of downregulated and upregulated DEGs comparing KOLF 56Q vs 21Q. **B.** Volcano plot of downregulated and upregulated DEGs comparing KOLF 79Q vs 21Q. **C.** Volcano plot of downregulated and upregulated DEGs comparing KOLF *HTT*KO vs 21Q. **D.** DEGs that contribute to pathway dysregulation in CREB Signaling in Neurons. **E.** DEGs that contribute to pathway dysregulation in G-Protein Coupled Receptor Signaling. **F.** DEGs that contribute to pathway dysregulation in FAK Signaling.

**Table S1:**
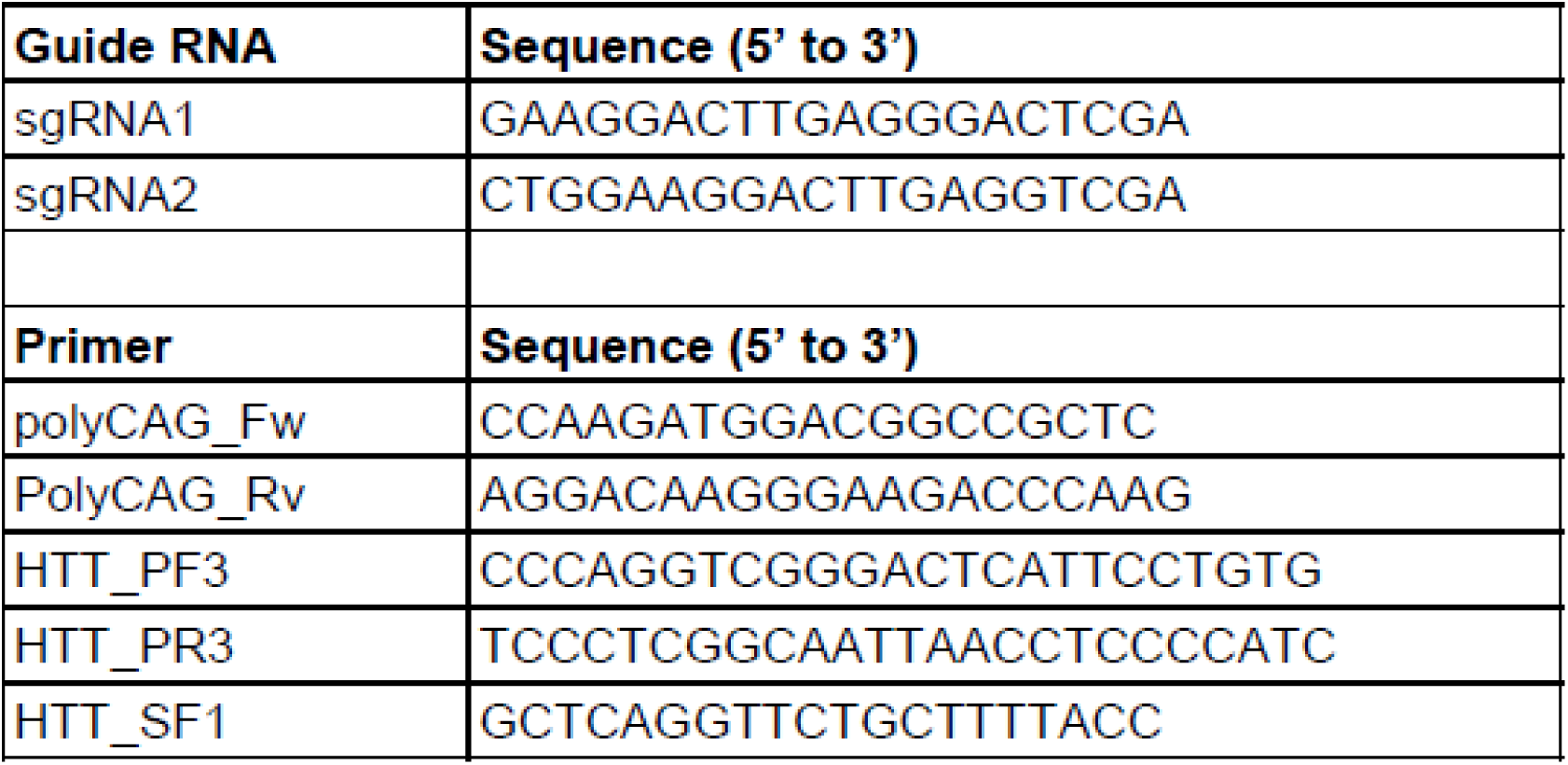
sgRNA and primer sequences used for the generation of the HTT isogenic series.

**Table S2:**
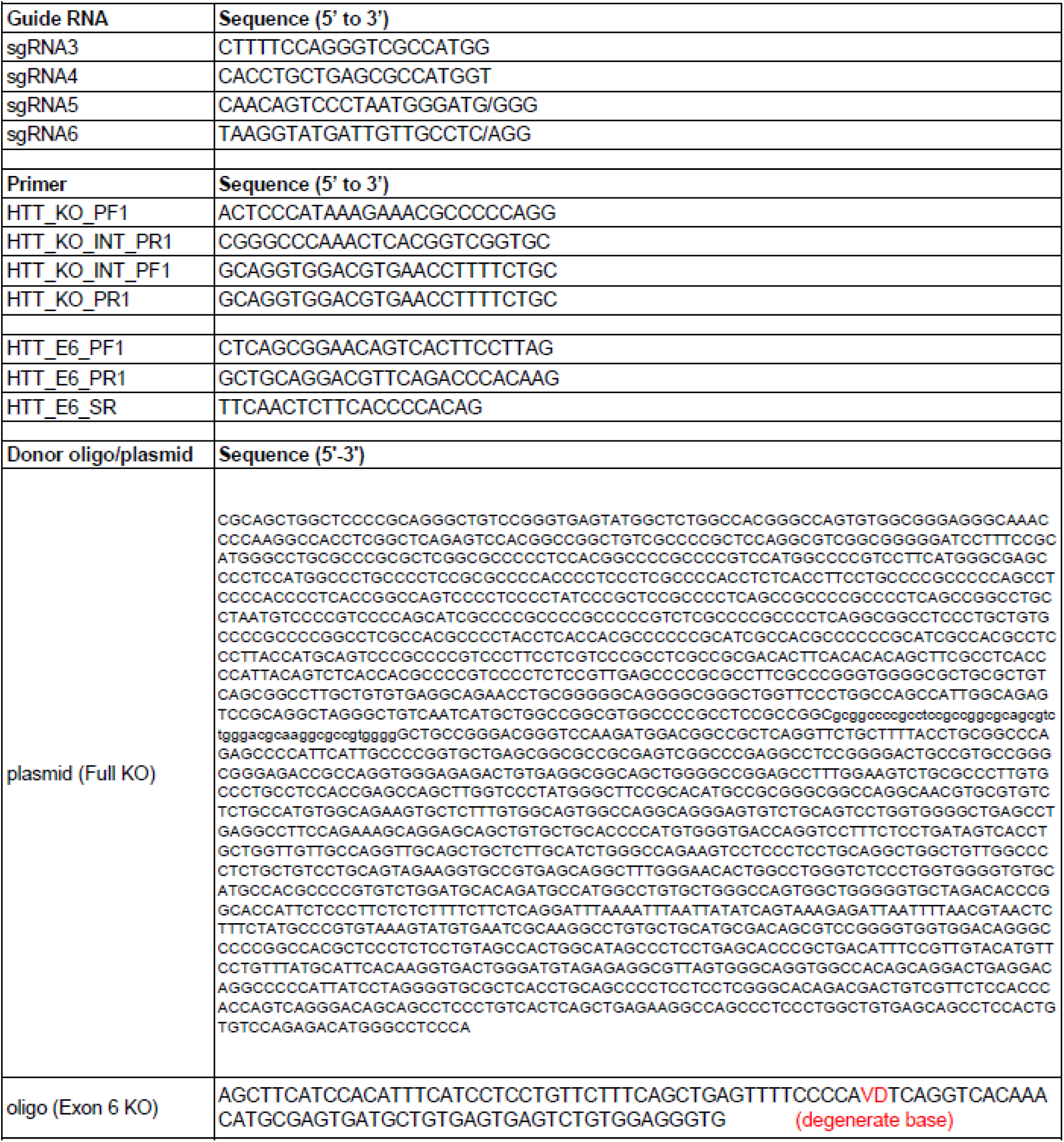
Sequences used for the generation of the HTT knock out lines.

**Table S3:**
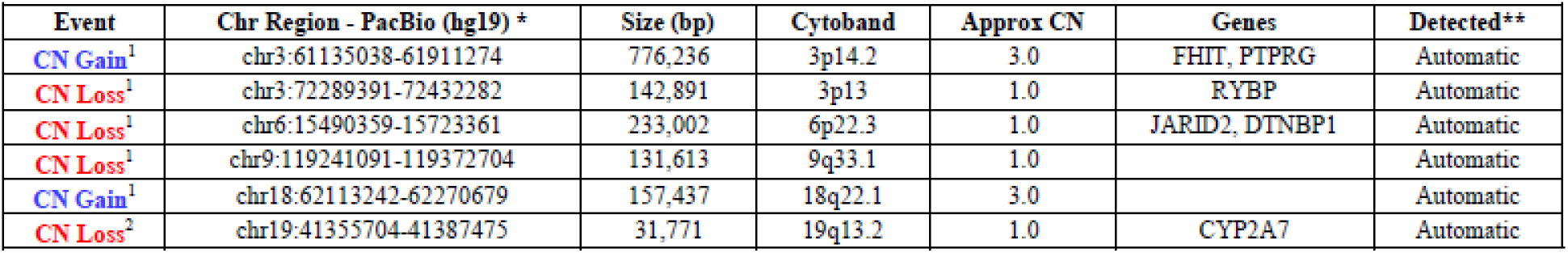
Known KOLF2-C1 CNVs. CNVs that have been identified in the KOLF2-C1 iPSC line, which is the original line from which KOLF2.1J was generated. These CNVs are present in all genetically engineered lines derived from KOLF2-C1 and KOLF2.1J. *CNVs were confirmed with PacBio Long Read sequencing data. CN Losses were identified with pav (doi:10.1126/science.abf7117, PMID: 33632895). CN Gains were identified with pbsv (https://github.com/PacificBiosciences/pbsv). **Known KOLF2-C1 CN events are automatically detected by VIATM in >90% of samples (Automatic). 1: CN events as reported by Garcia-Diaz et al^30^ and as confirmed by Ryan et al^31^ 2: CN event identified by C. Beck, unpublished analysis of PacBio WGS.

**Table S4:** Differentially expressed genes comparing 56Q vs 21Q, 79Q vs 21Q and KO vs 21Q MSNs.

## REFERENCES

1. MacDonald ME, Ambrose CM, Duyao MP, et al. A novel gene containing a trinucleotide repeat that is expanded and unstable on Huntington’s disease chromosomes. The Huntington’s Disease Collaborative Research Group. Cell 1993; 72: 971–983. DOI: 10.1016/0092-8674(93)90585-e.

2. Genetic Modifiers of Huntington’s Disease C. Identification of Genetic Factors that Modify Clinical Onset of Huntington’s Disease. Cell 2015; 162: 516–526. DOI:10.1016/j.cell.2015.07.003.

3. Genetic Modifiers of Huntington’s Disease Consortium. Electronic address ghmhe and Genetic Modifiers of Huntington’s Disease C. CAG Repeat Not Polyglutamine Length Determines Timing of Huntington’s Disease Onset. Cell 2019; 178: 887–900 e814. DOI: 10.1016/j.cell.2019.06.036.

4. Wexler NS, Lorimer J, Porter J, et al. Venezuelan kindreds reveal that genetic and environmental factors modulate Huntington’s disease age of onset. Proc Natl Acad Sci U S A 2004; 101: 3498–3503. 20040301. DOI: 10.1073/pnas.0308679101.

5. Walker FO. Huntington’s disease. Lancet 2007; 369: 218–228. DOI: 10.1016/S0140-6736(07)60111-1.

6. Takahashi K and Yamanaka S. Induction of pluripotent stem cells from mouse embryonic and adult fibroblast cultures by defined factors. Cell 2006; 126: 663–676. 20060810. DOI: 10.1016/j.cell.2006.07.024.

7. An MC, Zhang N, Scott G, et al. Genetic correction of Huntington’s disease phenotypes in induced pluripotent stem cells. Cell Stem Cell 2012; 11: 253–263. 20120628. DOI: 10.1016/j.stem.2012.04.026.

8. Xu X, Tay Y, Sim B, et al. Reversal of Phenotypic Abnormalities by CRISPR/Cas9-Mediated Gene Correction in Huntington Disease Patient-Derived Induced Pluripotent Stem Cells. Stem Cell Reports 2017; 8: 619–633. 20170223. DOI: 10.1016/j.stemcr.2017.01.022.

9. Tshilenge KT, Aguirre CG, Bons J, et al. Proteomic Analysis of Huntington’s Disease Medium Spiny Neurons Identifies Alterations in Lipid Droplets. Mol Cell Proteomics 2023; 22: 100534. 20230322. DOI: 10.1016/j.mcpro.2023.100534.

10. Malankhanova T, Suldina L, Grigor’eva E, et al. A Human Induced Pluripotent Stem Cell-Derived Isogenic Model of Huntington’s Disease Based on Neuronal Cells Has Several Relevant Phenotypic Abnormalities. J Pers Med 2020; 10 20201109. DOI: 10.3390/jpm10040215.

11. Ooi J, Langley SR, Xu X, et al. Unbiased Profiling of Isogenic Huntington Disease hPSC-Derived CNS and Peripheral Cells Reveals Strong Cell-Type Specificity of CAG Length Effects. Cell Rep 2019; 26: 2494–2508 e2497. DOI: 10.1016/j.celrep.2019.02.008.

12. Ruzo A, Croft GF, Metzger JJ, et al. Chromosomal instability during neurogenesis in Huntington’s disease. Development 2018; 145 20180129. DOI: 10.1242/dev.156844.

13. Conforti P, Besusso D, Brocchetti S, et al. RUES2 hESCs exhibit MGE-biased neuronal differentiation and muHTT-dependent defective specification hinting at SP1. Neurobiol Dis 2020; 146: 105140. 20201013. DOI: 10.1016/j.nbd.2020.105140.

14. Galimberti M, Nucera MR, Bocchi VD, et al. Huntington’s disease cellular phenotypes are rescued non-cell autonomously by healthy cells in mosaic telencephalic organoids. Nat Commun 2024; 15: 6534. 20240802. DOI: 10.1038/s41467-024-50877-x.

15. Stocksdale JT, Leventhal MJ, Lam S, et al. Intersecting impact of CAG repeat and huntingtin knockout in stem cell-derived cortical neurons. Neurobiol Dis 2025; 210: 106914. 20250419. DOI: 10.1016/j.nbd.2025.106914.

16. O’Regan GC, Farag SH, Casey CS, et al. Human Huntington’s disease pluripotent stem cell-derived microglia develop normally but are abnormally hyper-reactive and release elevated levels of reactive oxygen species. J Neuroinflammation 2021; 18: 94. 20210419. DOI: 10.1186/s12974-021-02147-6.

17. Ramos DM, Skarnes WC, Singleton AB, et al. Tackling neurodegenerative diseases with genomic engineering: A new stem cell initiative from the NIH. Neuron 2021; 109: 1080–1083. DOI: 10.1016/j.neuron.2021.03.022.

18. Pantazis CB, Yang A, Lara E, et al. A reference human induced pluripotent stem cell line for large-scale collaborative studies. Cell Stem Cell 2022; 29: 1685–1702 e1622. DOI: 10.1016/j.stem.2022.11.004.

19. Skarnes WC, Pellegrino E and McDonough JA. Improving homology-directed repair efficiency in human stem cells. Methods 2019; 164-165: 18–28. 20190616. DOI: 10.1016/j.ymeth.2019.06.016.

20. Fernandopulle MS, Prestil R, Grunseich C, et al. Transcription Factor-Mediated Differentiation of Human iPSCs into Neurons. Curr Protoc Cell Biol 2018; 79: e51. 20180518. DOI: 10.1002/cpcb.51.

21. Chen Y, Tristan CA, Chen L, et al. A versatile polypharmacology platform promotes cytoprotection and viability of human pluripotent and differentiated cells. Nat Methods 2021; 18: 528–541. 20210503. DOI: 10.1038/s41592-021-01126-2.

22. Reyes-Ortiz AM, Abud EM, Burns MS, et al. Single-nuclei transcriptome analysis of Huntington disease iPSC and mouse astrocytes implicates maturation and functional deficits. iScience 2023; 26: 105732. 20221206. DOI: 10.1016/j.isci.2022.105732.

23. Ferreira TA, Blackman AV, Oyrer J, et al. Neuronal morphometry directly from bitmap images. Nat Methods 2014; 11: 982–984. DOI: 10.1038/nmeth.3125.

24. McQuade A, Coburn M, Tu CH, et al. Development and validation of a simplified method to generate human microglia from pluripotent stem cells. Mol Neurodegener 2018; 13: 67. 20181222. DOI: 10.1186/s13024-018-0297-x.

25. Smith-Geater C, Hernandez SJ, Lim RG, et al. Aberrant Development Corrected in Adult-Onset Huntington’s Disease iPSC-Derived Neuronal Cultures via WNT Signaling Modulation. Stem Cell Reports 2020; 14: 406–419. 20200227. DOI: 10.1016/j.stemcr.2020.01.015.

26. Bolger AM, Lohse M and Usadel B. Trimmomatic: a flexible trimmer for Illumina sequence data. Bioinformatics 2014; 30: 2114–2120. 20140401. DOI: 10.1093/bioinformatics/btu170.

27. Anders S, Pyl PT and Huber W. HTSeq--a Python framework to work with high-throughput sequencing data. Bioinformatics 2015; 31: 166–169. 20140925. DOI: 10.1093/bioinformatics/btu638.

28. Love MI, Huber W and Anders S. Moderated estimation of fold change and dispersion for RNA-seq data with DESeq2. Genome Biol 2014; 15: 550. DOI: 10.1186/s13059-014-0550-8.

29. Kramer A, Green J, Pollard J, Jr., et al. Causal analysis approaches in Ingenuity Pathway Analysis. Bioinformatics 2014; 30: 523–530. 20131213. DOI: 10.1093/bioinformatics/btt703.

30. Gracia-Diaz C, Perdomo JE, Khan ME, et al. KOLF2.1J iPSCs carry CNVs associated with neurodevelopmental disorders. Cell Stem Cell 2024; 31: 288–289. DOI: 10.1016/j.stem.2024.02.007.

31. Ryan M, McDonough JA, Ward ME, et al. Large structural variants in KOLF2.1J are unlikely to compromise neurological disease modeling. Cell Stem Cell 2024; 31: 290–291. DOI: 10.1016/j.stem.2024.02.006.

32. Yusa K, Zhou L, Li MA, et al. A hyperactive piggyBac transposase for mammalian applications. Proc Natl Acad Sci U S A 2011; 108: 1531–1536. 20110104. DOI: 10.1073/pnas.1008322108.

33. Khakh BS, Beaumont V, Cachope R, et al. Unravelling and Exploiting Astrocyte Dysfunction in Huntington’s Disease. Trends Neurosci 2017; 40: 422–437. 20170531. DOI: 10.1016/j.tins.2017.05.002.

34. Khakh BS and Goldman SA. Astrocytic contributions to Huntington’s disease pathophysiology. Ann N Y Acad Sci 2023; 1522: 42–59. 20230302. DOI: 10.1111/nyas.14977.

35. Chandrasekaran A, Avci HX, Leist M, et al. Astrocyte Differentiation of Human Pluripotent Stem Cells: New Tools for Neurological Disorder Research. Front Cell Neurosci 2016; 10: 215. 20160926. DOI: 10.3389/fncel.2016.00215.

36. Stoberl N, Donaldson J, Binda CS, et al. Mutant huntingtin confers cell-autonomous phenotypes on Huntington’s disease iPSC-derived microglia. Sci Rep 2023; 13: 20477. 20231122. DOI: 10.1038/s41598-023-46852-z.

37. Choi YS, Lee B, Cho HY, et al. CREB is a key regulator of striatal vulnerability in chemical and genetic models of Huntington’s disease. Neurobiol Dis 2009; 36: 259–268. 20090724. DOI: 10.1016/j.nbd.2009.07.014.

38. Steffan JS, Kazantsev A, Spasic-Boskovic O, et al. The Huntington’s disease protein interacts with p53 and CREB-binding protein and represses transcription. Proc Natl Acad Sci U S A 2000; 97: 6763–6768. DOI: 10.1073/pnas.100110097.

39. Dowie MJ, Scotter EL, Molinari E, et al. The therapeutic potential of G-protein coupled receptors in Huntington’s disease. Pharmacol Ther 2010; 128: 305–323. 20100811. DOI: 10.1016/j.pharmthera.2010.07.008.

40. Lee HN, Hyeon SJ, Kim H, et al. Decreased FAK activity and focal adhesion dynamics impair proper neurite formation of medium spiny neurons in Huntington’s disease. Acta Neuropathol 2022; 144: 521–536. 20220720. DOI: 10.1007/s00401-022-02462-z.

41. Ratie L and Humbert S. A developmental component to Huntington’s disease. Rev Neurol (Paris) 2024; 180: 357–362. 20240412. DOI: 10.1016/j.neurol.2024.04.001.

42. Wiatr K, Szlachcic WJ, Trzeciak M, et al. Huntington Disease as a Neurodevelopmental Disorder and Early Signs of the Disease in Stem Cells. Mol Neurobiol 2018; 55: 3351–3371. 20170511. DOI: 10.1007/s12035-017-0477-7.

43. Sapp E, Kegel KB, Aronin N, et al. Early and progressive accumulation of reactive microglia in the Huntington disease brain. J Neuropathol Exp Neurol 2001; 60: 161–172. DOI: 10.1093/jnen/60.2.161.

44. Tai YF, Pavese N, Gerhard A, et al. Microglial activation in presymptomatic Huntington’s disease gene carriers. Brain 2007; 130: 1759–1766. 20070330. DOI: 10.1093/brain/awm044.

45. Crapser JD, Ochaba J, Soni N, et al. Microglial depletion prevents extracellular matrix changes and striatal volume reduction in a model of Huntington’s disease. Brain 2020; 143: 266–288. DOI: 10.1093/brain/awz363.

46. Crotti A, Benner C, Kerman BE, et al. Mutant Huntingtin promotes autonomous microglia activation via myeloid lineage-determining factors. Nat Neurosci 2014; 17: 513–521. 20140302. DOI: 10.1038/nn.3668.

47. Han I, You Y, Kordower JH, et al. Differential vulnerability of neurons in Huntington’s disease: the role of cell type-specific features. J Neurochem 2010; 113: 1073–1091. 20100317. DOI: 10.1111/j.1471-4159.2010.06672.x.

48. Reiner A, Dragatsis I, Zeitlin S, et al. Wild-type huntingtin plays a role in brain development and neuronal survival. Mol Neurobiol 2003; 28: 259–276. DOI: 10.1385/MN:28:3:259.

